# AI-based antibody discovery platform identifies novel, diverse and pharmacologically active therapeutic antibodies against multiple SARS-CoV-2 strains

**DOI:** 10.1101/2023.08.21.554197

**Authors:** Cristina Moldovan Loomis, Thomas Lahlali, Danielle Van Citters, Megan Sprague, Gregory Neveu, Laurence Somody, Christine C. Siska, Derrick Deming, Andrew J. Asakawa, Tileli Amimeur, Jeremy M. Shaver, Caroline Carbonelle, Randal R. Ketchem, Antoine Alam, Rutilio H. Clark

## Abstract

A critical aspect of a successful pandemic response is expedient antibody discovery, manufacturing and deployment of effective lifesaving treatments to patients around the world. However, typical drug discovery and development is a lengthy multi-step process that must align drug efficacy with multiple developability criteria and can take years to complete. In this context, artificial intelligence (AI), and especially machine learning (ML), have great potential to accelerate and improve the optimization of therapeutics, increasing their activity and safety as well as decreasing their development time and manufacturing costs. Here we present a novel, cost-effective and accelerated approach to therapeutic antibody discovery, that couples AI-designed human antibody libraries, biased for improved developability attributes with high throughput and sensitive screening technologies. The applicability of our platform for effective therapeutic antibody discovery is demonstrated here with the identification of a panel of human monoclonal antibodies that are novel, diverse and pharmacologically active. These first-generation antibodies, without the need for affinity maturation, bind to the SARS-CoV-2 spike protein with therapeutically-relevant specificity and affinity and display neutralization of SARS-CoV-2 viral infectivity across multiple strains. Altogether, this platform is well suited for rapid response to infectious threats, such as pandemic response.

**IMPORTANCE:** Expedient discovery and manufacturing of lifesaving therapeutics is critical for pandemic response. The recent COVID pandemic has highlighted the current inefficiencies and the need for improvements. To this end, we present our therapeutic antibody discovery platform that couples artificial intelligence (AI) and innovative high throughput technologies, and we demonstrate its applicability to rapid response. This platform enabled the isolation, characterization, and rapid identification of effective broadly neutralizing SARS-CoV-2 antibodies with good developability attributes, anticipated to fit our current process development and manufacturing platform. As such, this would benefit cost-of-goods and improve therapeutic access to patients. The AI-derived antibodies represent an advantageous therapeutic modality that can be developed and deployed fast, thus well suited for rapid response to infectious threats, such as pandemic response.

## INTRODUCTION

Antibodies represent an important class of biologics-based therapeutics with key benefits such as high specificity and affinity, longer-acting pharmacokinetics and superior safety profiles compared to small molecules (1, 2). Thus, antibody therapeutics are the fastest growing class of drugs on the market used for treatment of a wide range of human diseases, such as cancer, autoimmune, inflammatory, neural, metabolic, and infectious diseases.

The high potential of antibody therapeutics is, however, hampered by lengthy and expensive discovery and development processes. Indeed, candidate antibody therapeutics must undergo a complex multi-objective process and satisfy multiple criteria. This includes activity and specificity against a target, good pharmacokinetic and safety profiles, as well as suitable biophysical and manufacturing properties.

Traditionally, this is achieved in a sequential, funnel-like manner that lasts 10-15 years and costs approximately 2 billion USD (2). In particular, the recent COVID pandemic has highlighted the current inefficiencies and the need to provide affordable, high-quality treatment to patients around the globe and in much shorter timescales.

Progress in computational methods, technology and automation and their increasing integration in multiple aspects of the biopharmaceutical pipeline have the potential to revolutionize therapeutic antibody discovery and development. We have employed machine learning (ML) to generate novel, humanoid antibody sequences that both represent natural repertoires and are biased towards desirable developability features. To enable properties such as broad target and epitope engagement, focused efficacy, and suitable developability, we have developed an antibody Generative Adversarial Network (GAN), a new *in silico* engineering approach for designing a novel class of diverse, hyper-realistic antibodies, termed “humanoid” antibodies (3). The algorithm uses a modified Wasserstein-GAN for both single-chain (light or heavy chain) and paired-chain (light and heavy chain) antibody sequence generation (4). These GANs allow us to encode for antibodies with key properties of interest to create feature-biased libraries as the central benefit of our display libraries and to inform antibody engineering research. Our antibody GAN architecture 1) captures the complexity of the entire variable region of the standard human antibody sequence space, 2) provides a basis for rationally generating novel antibodies that span a larger sequence diversity than is explored by standard *in silico* generative approaches, and 3) provides, through transfer learning, an inherent method to bias the physical properties of the generated antibodies toward desired features which could lead to improved efficacy as well as chemical and biophysical properties which are critical for developability.

Here we present the construction and utilization of a GAN-generated phage display library of approximately 1 billion Fab antibodies. The Just Humanoid Antibody Library (J.HAL^®^) was successfully screened to isolate a panel of novel, diverse and pharmacologically active human monoclonal antibodies against SARS-CoV-2. These first-generation antibodies, without the need of affinity maturation, bind to the SARS-CoV-2 spike protein with therapeutically-relevant specificity and affinity, block the spike:human ACE2 receptor interaction and neutralize SARS-CoV-2 viral infectivity across several strains, as well as display desirable developability attributes.

To maximize efficiency of therapeutic discovery, we have paired the GAN-generated humanoid library with high throughput and sensitive technologies, such as next-generation sequencing, batch cloning, full-length IgG expression in 96 well blocks and a high throughput AlphaLISA-based binding screen.

The integration of automation and streamlined experimental and analysis workflows allows for the rapid screening, isolation and characterization of suitable IgG therapeutics. Our rapid response antibody discovery platform for addressing pathogenic disease demonstrates the promise of providing affordable, high-quality treatments to patients around the globe.

## MATERIALS & METHODS

### Phage library construction

Phagemids pADL-10b and pADL-20c (Antibody Design Labs, San Diego, CA) were used for construction of the GAN discovery library and were modified for expression of Fab antibody fragments as N-terminal pIII fusion proteins in *Escherichia coli.* Pools of synthetic gene fragments encoding variable heavy (VH) or variable light (VL) chains for each germline family were obtained from Integrated DNA Technologies (Coralville, IA). These fragments were flanked by >20 base pairs of sequence that included restriction sites used for cloning into the phagemid backbones.

A two-step cloning process was employed for combinatorial assembly of the V-regions into each phagemid. First, 50 ng of each of the 5 variable heavy chain pools was digested with NcoI/NheI and ligated with 300 ng of each phagemid backbone. The ligation reactions were cleaned up and concentrated using QIAquick PCR Purification Kit (Qiagen, Hilden, Germany) following the manufacturer’s protocol. Two microliters of this reaction were transformed into *E. coli* One Shot™ TOP10 electrocompetent cells (Thermo Fisher Scientific, Waltham, MA) in a 0.1 cm cuvette using a BTX ECM 630 electroporator (BTX, Holliston, MA) with the following settings: 1.8 kV, 600 Ohms, 25 µF. The transformants were recovered in 0.5 ml SOC media for 1 hour. Cells from the transformation reaction were then used to inoculate 150 ml TB supplemented with 50 µg/ml carb and 2% glucose and incubated at 30°C and 250 rpm overnight. Cultures were harvested the following day, and plasmid DNA was isolated using GenElute™ HP Maxiprep Kit (Millipore Sigma) following the manufacturer’s protocol.

Second, 5 µg of phagemid DNA harboring each variable heavy chain pool was digested using BsiWI/RsrII (for kappa chains) or BsiWI/KasI (for lambda chains) and ligated with 0.5 µg of each of the 5 VL chain pools that had been digested with the appropriate enzymes. The ligation reactions were cleaned up and concentrated using QIAquick PCR Purification Kit following the manufacturer’s protocol. Approximately two microliters of this reaction were transformed into MegaX DH10B T1^R^ Electrocomp™ cells (Thermo Fisher Scientific) in a 0.1 cm cuvette with the following settings: 2.0 kV, 200 Ohms, 25 µF. As many as 3-8 replicate transformations were performed for each germline pair to ensure the number of transformants exceeded the theoretical diversity by at least 10-fold. The transformants were recovered in 1 ml Recovery Media for 1 hour before pooling replicates and removing 10 µl for dilution plating on 2xYT plates supplemented with 100 µg/ml carb and 2% glucose, followed by overnight incubation at 30°C. The resulting colonies were used for estimating library size and for sequencing the VH and VL chains via colony PCR. The remainder of the transformation was used to inoculate 150 ml TB supplemented with 50 µg/ml carb and 2% glucose followed by overnight incubation at 30°C and 250 rpm. Cultures were harvested the following day and plasmid DNA was isolated using GenElute™ HP Maxiprep Kit, following the manufacturer’s protocol.

A total of 50 phagemid DNA preps were synthesized in this manner to complete the library (25 sub-libraries per each of 2 phagemid backbones). Colony PCR and Sanger sequencing were performed on a minimum of 24 clones per germline pair to assess correct insertion rate of the VH and VL domains.

### Phage library production

*E. coli* SS320 host cells (Lucigen Corporation, Middleton, WI) were electroporated using 30-50 ng of each sub-library DNA in a 0.1 cm cuvette with the following settings: 1.8 kV, 600 Ohms, 25 µF. Duplicate transformations were performed for each germline pair, recovered in 1 ml Recovery Media, and pooled before inoculating 2xYT broth containing 50 µg/ml carbenicillin and 2% glucose at an OD_600_ of 0.07. The cultures were then incubated with shaking at 250 rpm and 37°C until an OD_600_ ∼ 0.5, at which time the cells were infected with M13KO7 helper phage (Antibody Design Labs) at a multiplicity of infection (MOI) 25. The cells were continued incubating at 37°C without shaking for 30 minutes, followed by shaking at 200 rpm for 30 minutes. Cultures were centrifuged followed by medium replacement in 2xYT supplemented with 50 µg/ml carbenicillin and 25 µg/ml kanamycin. After overnight incubation at 30°C and 200 rpm, the phage particles were purified and concentrated by PEG/NaCl precipitation and resuspended in PBS containing 0.5% BSA and 0.05% Tween-20. Phage concentration was determined using a spectrophotometer, assuming 1 unit at OD_268_ is equivalent to 5×10^12^ phage/ml.

### Polyclonal phage ELISA (Fab display)

The display level of Fabs on phage was assessed for each sub-library using polyclonal phage ELISA. Briefly, 96-well MaxiSorp® assay plates (Nunc) were coated overnight at 4°C with anti-human Fab (Millipore Sigma) diluted 1:500 in PBS, then blocked in PBS containing 1% BSA for 1 hour at room temperature (RT). Phage preparations from each germline pair were serially diluted 1:25 in 2% non-fat dry milk in PBS, added to the plates, and allowed to incubate for 1 hour at RT before captured virions were detected using a 1:5000 dilution of anti-M13-HRP (Santa Cruz Biotechnology, Dallas, TX) for 1 hour at RT. All interval plate washes were performed 3 times in PBST (PBS supplemented with 0.1% v/v Tween-20). ELISAs were developed by addition of TMB solution (Thermo Fisher Scientific) and quenched using 10% phosphoric acid. Absorbance was read at A450 nm. Phage preparations derived from two commercial antibodies were used as high- and low-display Fab on phage controls (M-6240 and M-6239), respectively.

### Biopanning and clone sequencing

Biopanning was performed independently on two different antigens to isolate SARS-CoV-2 binders from the phage libraries. RBD (Cat. No. 40592-VNAH) and Spike S1 B.1.1.7 variant (Cat. no. 40591-V08H12) were acquired from Sino Biological (Beijing, China) and biotinylated using EZ-Link™ Sulfo-NHS-Biotin kit (Thermo Fisher Scientific) as recommended by the manufacturer. In the first round of selection, approximately 1×10^13^ phage particles were incubated in 1 ml SuperBlock™ (Thermo Fisher Scientific) for 1 hour. The blocked phages were next incubated with 100 nM biotinylated antigen for 1 hour, followed by incubation with magnetic streptavidin beads (Dynabeads™ M-280 Streptavidin, Thermo Fisher Scientific) for 15 minutes. Panning was performed by washing the beads 2 times with PBS-T (PBS with 0.1% Tween-20) for 5 seconds each, followed by two additional washes in PBS-LT (PBS with 0.01% Tween-20) for 5 seconds each. The bound phages were eluted from the beads using 0.2M glycine (pH 2.5) for 10 minutes and neutralized with 1M Tris-HCl (pH 8.0). All incubations were performed using the KingFisher™ Flex at room temperature. Eluted phages were used to infect E. coli ER2738 cells (Lucigen Corporation) for titer determination and overnight amplification. Prior to the next round, the amplified phage was precipitated in 20% PEG/2.5M NaCl for at least 20 minutes on ice. Subsequent rounds of panning were performed similarly but had reduced input phage (5×10^12^ pfu), reduced antigen concentrations (2^nd^ round: 25 nM and 3^rd^ round: 10 nM), and longer wash times (2^nd^ round: 1.5 minutes and 3^rd^ round: 15 minutes) for increased stringency. Following the third round of panning, single clones were picked at random for colony PCR and Sanger sequencing of VH and VL chains. Unique sequences were identified for assessment of RBD or S1 B.1.1.7 binding using monoclonal phage ELISA.

### Next-generation sequencing and PCR recovery of enriched Fabs

The VH domain of the third-round RBD panning outputs was deep sequenced to enable targeted recovery of low-frequency Fabs using PCR. First, phage particles eluted off beads were used directly as template in PCR to amplify the VH region. These primers contained Read 1 and Read 2 adapters for paired-end sequencing on the Illumina platform. PCR products were column purified using the QIAquick PCR Purification Kit (Qiagen) and sent to Genewiz for sequencing using their Amplicon-EZ service. Paired-end sequences were merged using Geneious Prime software, followed by sequence alignment using internally developed proprietary software (Abacus). Sequences were ranked in descending order by count, and VH sequences with a frequency of at least 0.01% in the panning output were identified. In total, 164 VH sequences that had not previously been sampled in earlier screens were targeted for recovery by PCR. Next, reverse PCR primers were designed for each targeted VH sequence such that the 5’ end annealed in the CH1 domain and the 3’ end anchored in the unique HCDR3 region. Reverse primers were 72 bp in length and obtained from Integrated DNA Technologies in 11 distinct oligo pools (“oPools”). Each oPool was paired with a universal forward primer annealing in the signal peptide for light chain, enabling amplification of a DNA fragment containing VL, CL, and the targeted VH domains using phage panning eluates as a template. This fragment was gel purified, digested using appropriate restriction enzymes, and ligated back into the phagemid backbone. Sanger sequencing was performed on 384 picked colonies, and of these, 51 unique Fabs were selected for conversion to IgG using the methods described above.

### Monoclonal phage production

Single clones harboring a unique Fab fusion protein were inoculated into 150 µl 2xYT broth supplemented with 50 µg/ml carbenicillin, 15 µg/ml tetracycline, and 2% glucose and cultivated in 96 deep-well plates overnight at 37°C with rigorous shaking. Five µl of the overnight cultures was then transferred to new deep-well plates containing 100 µl 2xYT with 50 µg/ml carbenicillin, 15 µg/ml tetracycline, and 2% glucose and incubated at 37°C with rigorous shaking until an OD_600 nm_ ∼ 0.5. M13KO7 helper phage was added to each well at MOI 25, and plates were incubated without agitation at 37°C for 1 hour before medium replacement to 2xYT containing 50 µg/ml carbenicillin and 25 µg/ml kanamycin and overnight incubation with rigorous shaking at 30°C. Phage supernatants were harvested after centrifugation and diluted 1:3 in 2% non-fat dry milk in PBS for use in the monoclonal phage ELISA.

### Monoclonal phage ELISA (antigen binding)

Binding of enriched Fabs displayed on phage to recombinant antigen was determined using phage ELISA. Briefly, 96-well MaxiSorp^®^ assay plates (Nunc) were coated overnight at 4°C with 5 µg/ml NeutrAvidin (Thermo Fisher Scientific), then blocked in PBS containing 1% BSA for 1 hour at room temperature (RT). Biotinylated antigen (i.e., RBD or S1 B.1.1.7) was added at a concentration of 1 µg/ml for 30 minutes. Diluted phage preparations were added and allowed to incubate for 1 hour at RT before captured virions were detected using a 1:5000 dilution of anti-M13-HRP (Santa Cruz Biotechnology) for 1 hour at RT. All interval plate washes were performed 3 times in PBST (PBS supplemented with 0.1% v/v Tween-20). ELISAs were developed by addition of TMB solution (Thermo Fisher Scientific) and quenched using 10% phosphoric acid. Absorbance was read at A_450 nm_. Plate wells coated with an irrelevant antigen (biotinylated CD40, Sino Biological, Cat. No. 10774-H08H) were processed in parallel to assess specificity of binders. Phage supernatants from two known RBD binders were included as positive controls.

### Batch conversion of Fab to full-length IgG

After successful enrichment of SARS-CoV-2 binders was confirmed by phage ELISA, pooled panning outputs were subcloned into a mammalian expression vector in a manner that converts Fabs to full-length IgG while simultaneously maintaining VH-VL pairing for high throughput screening. First, phagemid DNA was purified from phage panning outputs as follows. ER2738 cells infected with pooled phage output after the second and third rounds of panning were plated densely on 2xYT agar supplemented with 100 µg/ml carbenicillin and 2% glucose and incubated overnight at 30°C. Cells were then scraped in 1-2 ml 2xYT broth containing 50 µg/ml carbenicillin and 2% glucose, and plasmid DNA from 200-500 µl of the cell suspension was purified using QIAprep Spin Miniprep Kit (Qiagen) following the manufacturer’s protocol. The process of converting phage panning outputs from Fabs to full-length IgG was then performed in two steps. First, a linear fragment containing VL and VH was isolated from the phagemid backbone using restriction digest with BsiWI and NheI. This fragment was ligated into the mammalian vector such that the VL is cloned downstream of a CMV promoter and mammalian signal peptide, and VH is cloned upstream of IgG1 (i.e., CH1, hinge, CH2, and CH3 domains). This intermediate plasmid thus consists of antibody cassettes with complete light chain (LC) and heavy chain (HC) IgG1 regions but lacks the HC control elements. In the second cloning step, plasmid DNA was purified from the intermediate pool and digested with RsrII or KasI (for kappa or lambda chains, respectively) and NcoI to replace the fragment between VL and VH with a fragment containing the constant kappa or lambda domain, polyA signal, and a CMV promoter and signal peptide for HC expression. The final plasmid product contains dual CMV expression cassettes for full-length IgG expression while maintaining the VH-VL pairing of the original selected Fab fragment. A total of 487 single colonies from the batch cloning were randomly picked and cultured overnight in 0.9 ml TB supplemented with 30 µg/ml kanamycin. Colony PCR and Sanger sequencing of VH and VL domains was performed on each clone. In addition, plasmid DNA was isolated from each clone using QIAprep 96 Turbo Kit (Qiagen) following the manufacturer’s protocol.

### Expression and quantitation of full-length IgG

The Expi293™ Expression System (Thermo Fisher Scientific) was utilized for transient production of full-length IgG from the 463 randomly picked sequences following batch reformatting. Expi293F cells were transfected using ExpiFectamine™ 293 reagent in 0.7 ml culture volumes with approximately 0.7 µg plasmid DNA per well, following the manufacturer’s protocol for 96-well microtiter plates. For a positive control, trastuzumab was cloned into the same mammalian expression vector and transfected in the same manner. Cells transfected with transfection reagents and no plasmid DNA served as negative controls. The transfected plates were covered with “System Duetz” sandwich covers (EnzyScreen BV, Netherlands), secured onto a 3 mm orbital platform shaker inside a humidified 37°C tissue culture incubator with 5% CO_2_, and shaken at 1000 rpm for 4 days. Enhancers were added 18-24 hours post-transfection. Culture supernatants were harvested by centrifuging the 96-well plates 3,000 rpm for 5 minutes to pellet cell debris. The cleared supernatants were diluted in fresh expression medium for titer quantification on the Octet® RED96 System (ForteBio Inc, Fremont, California) using Protein A biosensors. The sample IgG titers were determined by comparison to a previously established standard curve using known IgG concentrations and reported as µg/ml.

### AlphaLISA binding screen

An AlphaLISA^®^ (Perkin Elmer) bead-based luminescent amplification binding assay was performed to screen antibodies that specifically bound SARS-CoV-2 Spike protein and did not bind an irrelevant antigen. Briefly, 10 µl of unpurified transfection supernatant diluted 1:200 in 1x Immunoassay buffer (Perkin Elmer cat# AL000F) was pre-incubated with either 10 µl biotinylated SARS-CoV-2 spike protein (3 nM final concentration; R&D Systems cat# BT10549), or with 10 µl biotinylated irrelevant antigen (3 nM final concentration; R&D Systems cat# AVI10538-050) for 20 minutes at room temperature in 96 half area white microplates (Corning cat# CLS3693). Subsequently, 10 µl of Protein A AlphaLISA^®^ Acceptor beads (10 µg/ml final concentration; Perkin Elmer cat# AL101C) were added, and further incubated for 1 hour at room temperature in the dark with shaking (250 rpm). Following this incubation, 20 µl of Streptavidin Donor beads (40 µg/ml final concentration; Perkin Elmer cat# 6760002S) were added, and the microplates were further incubated for 1 hour at room temperature in the dark with shaking (250 rpm) and read on FLUOStar^®^ Omega multi-mode microplate reader instrument (BMG Labtech) (excitation 680 nm, emission 615 nm). Data were graphed using GraphPad Prism software.

For dose-dependent binding, positive unpurified supernatants were normalized to a starting concentration of 300 ng/ml (final concentration) and 9 point 1:3 serial dilutions were performed in 1x Immunoassay buffer prior to pre-incubation with biotinylated SARS-CoV-2 spike protein (3 nM final concentration) and an AlphaLISA^®^ assay was performed as described above.

### Inhibition of SARS-CoV-2 spike binding to human Angiotensin Converting Enzyme 2 (ACE-2)

Antibody supernatants that specifically bound SARS-CoV-2 spike protein were tested for their ability to block binding of SARS-CoV-2 spike protein to human ACE-2. Briefly, MaxiSorp^TM^ ELISA plates (Biolegend cat# 50-712-278) were coated with human ACE-2 Fc chimera protein (R&D Systems cat# 10544-ZN) in cold PBS (100 μl per well; 0.5 µg/ml) overnight at 4° C. The plates were then washed using PBS with 0.05% Tween 20 (wash buffer) and blocked with PBS, 1% (v/v) BSA for 1 hour at 37°C in a 5% CO_2_ incubator. Unpurified transient transfection supernatant samples (50 µl) were pre-incubated with biotinylated SARS-CoV-2 spike protein (R&D Systems cat# BT10549) (50 µl, 0.3 µg/ml final concentration in PBS with 1% BSA and 0.05% (v/v) Tween 20 (binding buffer)) for 20 minutes at room temperature, then transferred to the ELISA plates and incubated for 2 hours at room temperature with shaking (250 rpm). After this incubation, the plates were washed three times with wash buffer, and 100 µl Streptavidin - Horseradish Peroxidase (Thermo Scientific cat# N100) (1:5000 dilution in binding buffer) was added onto the wells and incubated for 45 minutes at room temperature. After a 3× wash step with wash buffer, the plates were developed using 3,3′,5,5′-Tetramethylbenzidine (TMB) substrate (100 µl; Pierce cat# 34022), and the reaction was quenched using Phosphoric Acid 10% (v/v) (100 µl; Ricca Chemical Company cat# 5850-16). The well absorbance was read at 450 nm on FLUOStar^®^ Omega multi-mode microplate reader instrument (BMG Labtech). Antibody blockade, represented as percent inhibition relative to the binding of spike protein alone, was calculated and graphed using GraphPad Prism software.

For dose-dependent inhibition, unpurified supernatants were normalized to a starting concentration of 30 µg/ml (final concentration) and 10 point 1:3 serial dilutions were performed in binding buffer prior to pre-incubation with biotinylated SARS-CoV-2 spike protein (0.3 µg/ml final concentration) and ELISA assay performed as described above.

### rVSV production

The pseudotyped ΔG-luciferase (G*ΔG-luciferase) rVSV was purchased from Kerafast (catalog number: EH1020-PM) and was used to produce VSV-ΔG pseudotyped with the SARS-CoV-2 spike protein. Briefly, HEK-293T cells were plated in T75 flask and incubated at 37°C, 5% CO2. Twenty-four hours post-seeding, cells were transfected with 12 µg of plasmid expressing the spike from various variants using Fugene-6 (Promega; catalog number: E2691) according to the manufacturer’s instructions and incubated at 37°C, 5% CO2. One hour later, cells were infected at a multiplicity of infection (MOI) of 5 with pseudotyped ΔG-luciferase (G*ΔG-luciferase) rVSV and incubated at 37°C, 5% CO2 overnight. In the morning, cells were washed 3 times with PBS, fresh media was added. Cells were incubated for approximately 24 hours at 37°C, 5% CO2, and the supernatants were then collected, clarified by low-speed centrifugation (1320×g for 10 min), and were aliquoted and stored at −80°C. The rVSVΔG-SARS-CoV-2-spike proteins were titered by TCID_50_ (50% tissue culture infectious dose). Briefly, pseudotyped virus titration is carried out by using 96-well plates. A549-hACE2/hTMPRSS2 cells were infected with 10-fold serial dilutions in 4 replicates. After infection with the pseudotyped virus at 37°C, and 5% CO2, for 24 hours, the luciferase substrate was added for chemiluminescence detection (Promega; Luciferase Assay system; catalog number: E1500). The detected raw data were used to calculate the TCID_50_ according to the Reed-Muench method. The positivity of the luciferase readout was set to 4 times the luminescence activity of the uninfected control.

### Spike variants used

The spike from the following variants of interest and concern were used to generate rVSVΔG-SARS-CoV-2-spike pseudotypes: SARS-CoV-2 USA-WA1/2020 (Lineage A) SARS-CoV-2 USA-WA1/2020, D614G (Lineage A), SARS-CoV-2 hCoV-19/England/204820464/2020, variant alpha (Lineage B.1.1.7), SARS-CoV-2 hCoV-19/South Africa/KRISP-K005325/2020, beta variant (Lineage B.1.351), SARS-CoV-2 hCoV-19/Japan/TY7-503/2021, gamma variant (Lineage P.1), SARS-CoV-2 hCoV-19/USA/PHC658/2021, delta variant (lineage B.1.617.2), SARS-CoV 2 hCoV-19/USA/ MD-HP20874/2021, omicron BA.1 variant (Lineage B.1.1.529.1), hCoV-19/South Africa/CERI-KRISP-K032307/2021, omicron BA.2 variant (Lineage B.1.1.529.2).

### Neutralization assay with rVSVΔG-SARS-CoV-2-Spike

For the neutralization procedure, the virus corresponding to a multiplicity of infection (MOI) of 0.1 was incubated with increasing concentrations (10-fold serial dilutions) of test antibodies for 1 hour at 37°C to allow the antibody to bind to the viral surface envelope glycoproteins. A549-hACE2/TMPRSS2 cells were then infected with the virus/antibody mixture and incubated for 20 hours at 37°C, 5% CO2. At 20 hours post-infection, the luciferase was detected using the Luciferase Assay system (Promega; catalog number: E1500) according to the manufacturer’s instructions. Briefly, the cells were washed once with PBS, lysed using 1x Passive Lysis Buffer for 30 minutes. The luciferase substrate was then added, and luminescence was measured using a 96-well plate reader (Envision microplate reader; Perkin Elmer) with an integration time of 0.1s per well.

The results were normalized to a percentage of neutralization (Neutralization %) where the cell control (non-infected condition) and the virus control (non-treated, infected condition) are used to set 100% neutralization and 0% neutralization, respectively.

### Neutralization assay with SARS-CoV-2 isolates

For the neutralization procedure, the virus at a MOI of 0.1 was incubated with increasing concentrations (10-fold serial dilutions) of S309 (11) or test antibodies for 1 hour at 37°C to allow the antibody to bind to the viral surface envelope glycoproteins. A549-hACE2/TMPRSS2 cells were then infected with the virus/antibody mixture and incubated for 40 hours at 37°C, 5% CO2. Remdesivir at 5 µM was also used as a control (Sigma-Aldrich, St. Louis, MO, USA). At 40 hours post-infection infectivity was assessed by quantifying intracellular viral genomes by RT-qPCR. Briefly, the supernatant was removed, and cells were washed once with PBS 1X. Cells were then lysed with 100 µL of lysis buffer. Total RNA was extracted using the SV 96 Total RNA Isolation System and Vac-Man 96 (Promega; catalog number: Z3500), according to the manufacturer’s instructions. Taqman Fast Virus 1-step kit (Thermo Fisher; catalog number 4444434) with oligos and probe specific to the N gene of SARS-CoV2 was used for SARS-CoV-2 RNA detection according to the manufacturer’s instructions. Standard range was prepared by 10-fold serial dilution of qPCR Control RNA from inactivated SARS-CoV-2. The QuantStudio 7 Flex cycler (Thermo Fisher) was used for amplification. The results were expressed as copies of viral genome equivalent per µl of RNA.

### SARS-CoV-2 isolates

Isolates USA-WA1/2020 (A), cat. no.NR-52281, hCoV19/USA/PHC658/2021 (Delta B.1.617.2), cat. no. NR-55611 and hCoV-19/USA/MD-HP20874/2021 (Omicron B.1.1.529, BA.1), cat. no. NR-56461 were obtained through BEI Resources, NIAID, NIH.

### Surface Plasmon Resonance (SPR) kinetics characterization

SPR was used to determine the affinities of selected candidates for spike proteins from multiple SARS-CoV-2 strains (Wuhan (lineage A), Denmark – Mink Cluster-5 (lineage B.1.1.298), Japan (or Brazil)(lineage P.1), USA/CA San Diego (lineage B.1.1.7), Indian-Kappa variant (lineage B.1.617.1), UK variant (lineage B.1.1.7), and South African variant (lineage B.1.351)). Briefly, kinetics measurement was performed by SPR using the Carterra LSA instrument (carterra-bio.com). The instrument preparation and operation were conducted as recommended by the manufacturer. All kinetics measurements were performed at 25°C. Purified anti-SARS-CoV-2 antibodies of the invention were printed for 10 minutes using 20 µg/ml material onto sensor chips conjugated with anti-human Fc capture antibody. Appropriate controls were printed alongside novel antibodies to confirm capture integrity of sensor chip and associated system validation. Purified spike protein material was serially diluted to 8 concentrations starting at 500 nM. Buffer blanks were passed over the sensor chip prior to spike protein injection for sensor array chip preparation and assess baseline signal level. Each spike protein kinetic series was injected from low to high concentration to generate eight sensorgrams per antibody:spike variant interaction. Replicates for each 8-sensorgram set were performed. The data were processed and analyzed by Carterra’s KIT software tool by interspot referencing and double referencing the data and then by fitting them to a simple Langmuir binding model using global k_a_, k_d_, and R_max_ values per spot. Fit quality was determined by inspection of the residuals.

### Biophysical characterization sample preparation

Samples were buffer exchanged against 10 diavolumes of 20mM sodium phosphate, 150mM sodium chloride, pH 7.1 (PBS) using a centrifugal filter with a 30 kDa molecular weight cut off (Amicon). After buffer exchange, samples were normalized to 1 mg/ml using a Lunatic protein concentration plate format instrument (Unchained Labs).

### Differential Scanning Fluorimetry

Thermal transition temperature(s) and weighted shoulder scores were determined by DSF according to the method previously described (5). A single biological sample was divided, and the assay ran twice per molecule type. Additional information was also obtained from a parameter termed the weighted shoulder score (WSS) which accounts for multiple pieces of information from the unfolding curve (6).

### Low pH stability

Stability during a low pH hold was determined according to the method previously described (7). The increase in high molecular weight of the low pH exposed sample as compared with the control sample is reported.

### Chemical unfolding

The chemical unfolding assay was completed as previously described (8) with some modifications. After a 3-day incubation in 32 guanidine hydrochloride (GND) concentrations, the samples were measured on a Fluorescence Innovations SUPR-UV plate reader (excitation: 275 nm, emission: 300-440 nm). The measured fluorescence intensity at 362 nm was corrected for scattering and stray light, the unfolding curve was generated by graphing each corrected intensity against the GND concentration and the inflection point is reported.

### Relative solubility

Solubility was assessed according to the method previously described (5). Analysis was done in PBS buffer (20mM sodium phosphate and 150mM sodium chloride pH 7.1) and a final PEG 10,000 concentration ranging from 7.2% to 9.6%. Percent recovery relative to a 0% PEG control was determined and average recovery across the PEG concentration range is reported.

### Self-Interaction Nanoparticle Spectroscopy (SINS)

SINS measurements were performed according to the method previously described (9). Briefly, gold nanoparticles (Ted Pella) were conjugated overnight with an 80:20 ratio of anti-human and anti-goat antibodies (Jackson Immuno Research). Unreacted sites were blocked using an aqueous 0.01% (w/v) polysorbate 80 solution. Conjugated gold nanoparticles were then concentrated by centrifugation and removal of 95% of the supernatant. Analysis was carried out in PBS (20mM phosphate, 150mM NaCl, pH 7.1) at a protein concentration of 0.05 mg/ml reacted with 5 µl of concentrated conjugated gold nanoparticles. After a 2-hour incubation, absorbance spectrum from 400-600 nm was collected using a Spectrostar Nano plate reader at 2nm steps. The wavelength maximum of the spectrum peak is reported. Three assay replicates were made from each of the pooled expressed antibodies.

### Standup Monolayer Absorption Chromatography (SMAC)

SMAC measurements were performed according to the method previously described (10). Retention times were determined using a Dionex UPLC equipped with a Zenix column (Sepax Technologies) and a running buffer comprised of 150mM sodium phosphate pH 7.0.

## RESULTS

### J.HAL^®^ library construction

Two different phagemids were used for construction of the GAN discovery library. pADL-20c (Antibody Design Labs) is designed for high-level display of a gene of interest on the N-terminus of pIII under the control of the lac promoter. This phagemid was modified for bicistronic expression of Fab antibody fragments using the *E. coli* heat-stable enterotoxin (STII; for light chain) and pectate lysate (pelB; for heavy chain) signal sequences for periplasmic translocation of the Fab-fusion proteins. A hexahistidine tag and a FLAG tag were added to the C-terminus of the CH1 and light chain constant domains, respectively, and the amber stop codon upstream of gIII was removed to allow expression of the fusion protein in SS320 host cells. pADL-10b (Antibody Design Labs) is a low-display phagemid containing a copy of the lacI transcriptional repressor gene and a tHP transcriptional terminator upstream of the promoter. These elements are used for tighter repression of the lac operon during amplification of the library to prevent loss of clones bearing toxic Fab-pIII fusion proteins. It was modified in a similar manner as pADL-20c for Fab display, except that the pelB signal sequence was used for both heavy and light chain secretion. We hypothesized that binders would be easier to obtain from the pADL-20c phagemid due to higher display levels, but that the clonal distribution would be more normalized in the pADL-10b phagemid due to reduced effect of toxicity on growth rate. To increase the likelihood of isolating binders against the target of interest, the GAN antibody sequences were cloned into both phagemids independently, resulting in the production of two distinct libraries, each containing approximately 3.5 x 10^10^ transformants representing a theoretical diversity of 1 billion VH and VL pairs. Sanger sequencing of VH and VL from randomly picked colonies indicated that approximately 70% of clones in the library had correctly assembled pairs. The remaining 30% of clones had non-productive pairs, including stop codons, frameshifts, or truncated inserts. Fab-displaying phages were produced from each library. Phage ELISA was used to assess expression and display of the GAN-generated Fab fragments on phage (Figure 1). Display levels were estimated by capturing serial dilutions of purified phage on ELISA plates coated with anti-human Fab and detecting with anti-M13 antibodies conjugated to HRP. There is considerable variability in display levels among the various germline pairings, but in general, all pairs tend to fall between the high- and low-display control Fabs (M-6240 and M-6239, respectively). Furthermore, germline pairs with IGHV5-51, IGKV4-1, and IGLV1-40 appear to have lower display levels than other pairs in this library. Whether or not these differences are significant or represent differential binding of the anti-human Fab capture antibody used in the ELISA is an area of further investigation. Germline pairs cloned into the higher-display phagemid pADL-20c had similar trends as those in pADL-10b (data not shown).

**Figure 1.**
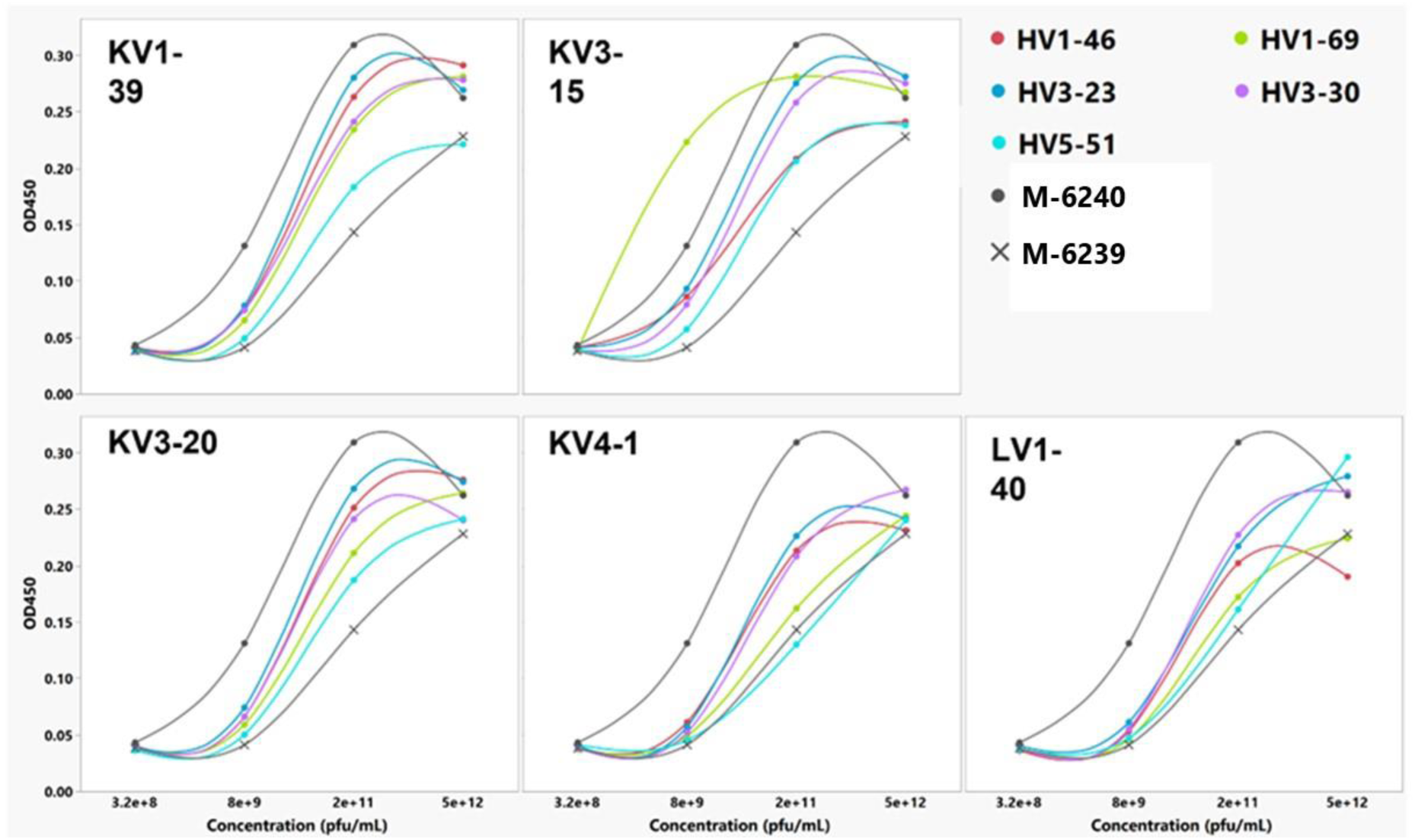
Fab phage display levels of 25 GAN sub-libraries. ELISA showing Fab expression of 25 GAN sub-libraries from polyclonal phage. Graphs are grouped by the LC germline family. High- and low-expressing Fab on phage controls (M-6240 and M-6239, respectively) are included on each graph as a reference.

### Biopanning and Isolation of anti-SARS-CoV-2 Fabs

Biopanning was performed independently on two different SARS-CoV-2 antigens (Wuhan RBD and B.1.1.7 S1 (Sino Biological cat# 40592-VNAH and 40591-V08H12, respectively)) to isolate SARS-CoV-2 binders from the phage libraries. Biopanning was performed with increasing stringency of selection at each round, ranging from 100nM to 10nM biotinylated antigen. Following the third round of panning, single clones were picked at random for colony PCR and Sanger sequencing of VH and VL chains. Unique sequences were identified for assessment of RBD or S1 B.1.1.7 binding using monoclonal phage ELISA.

In parallel, ELISA assays were performed on CD40, an irrelevant antigen, to verify the specificity of the binders for RBD. Duplicate clones harboring the same Fab sequence were included, if possible. A total of 22 unique Fab sequences were identified as binding specifically to RBD. This was defined as having OD_450_ values greater than twice the OD_450_ signal as compared to binding to CD40 (Figure 2) thus confirming the efficiency of the selection strategy. The two positive control Fabs included in the assay were verified to be specific for RBD binding. Based on these results, panning was terminated after the third round, and selected phage outputs were converted in batch to full-length IgG for high throughput screening and characterization.

**Figure 2.**
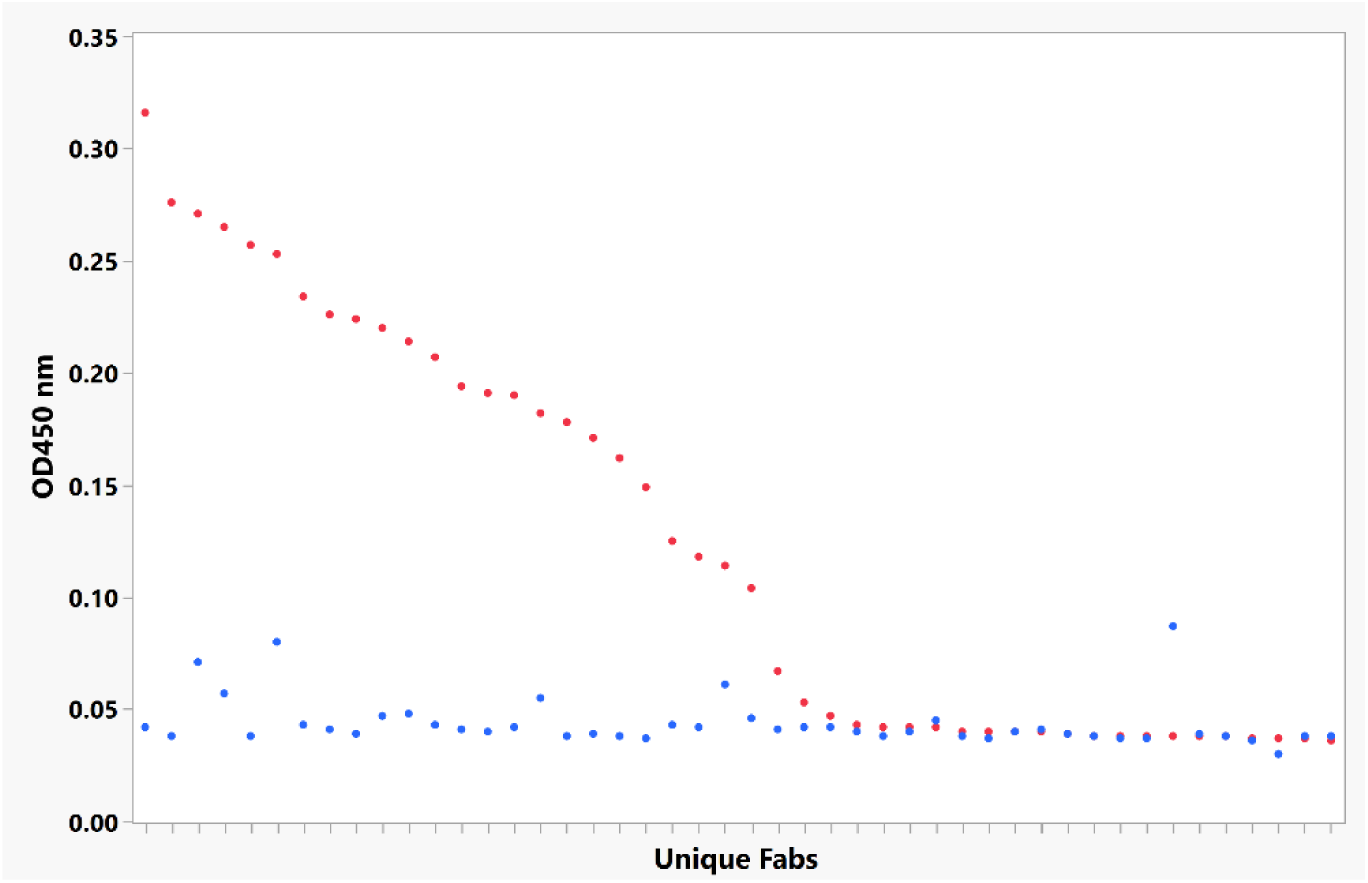
Isolated Fabs display specific binding to SARS-CoV-2 target antigen. Monoclonal phage ELISA analysis of unique Fab clones enriched after three rounds of selection against RBD. Binding to immobilized RBD (red) or the irrelevant antigen CD40 (blue) are shown. RBD specificity was defined as having RBD binding intensities at least twice that against CD40.

### Next-generation sequencing (NGS) and PCR recovery of enriched Fabs

To identify additional Fabs targeting the RBD of SARS-CoV-2, a high-throughput screen was implemented in parallel with random clone screening. The workflow used next generation sequencing coupled with PCR to recover less represented clones within the enriched third round outputs, which are not likely to be sampled by random colony picking. We first used PCR to amplify the VH region directly from the phage eluates after the third round of selection against RBD for analysis of clonal diversity by NGS. Based on these results, VH sequences with frequencies of 0.01% or higher that had not previously been tested by random screening were identified. We next designed pools of reverse PCR oligos that began in the CH1 domain and anchored in the unique HCDR3 region of each of those sequences. Using a universal forward primer that anneals in the signal peptide for light chain, we were able to recover linear fragments linking the targeted VH to its paired VL (VL-constant light chain-VH). The linear fragments were cloned back into the phagemid backbone, and 4 plates of single clones were picked for Sanger sequencing of the VH and VL domains. Of the 90 unique sequences identified in this panel, 68 were unique and had not been previously tested for binding to RBD. We selected 51 for IgG reformatting and transient mammalian expression.

### Fab to full-length IgG conversion and mammalian expression

We used a high throughput, two-step batch cloning method to convert enriched Fabs to full-length IgG, while simultaneously maintaining VH-VL linkage, for mammalian expression and further characterization. Following conversion of the Fabs to IgG, Sanger sequencing was performed on >400 picked colonies to obtain VH and VL sequences. Clones that contain the same sequence, as well as clones with non-productive sequences containing frameshifts, stop codons, or truncations, are removed. Unique productive sequences were then transiently expressed in a mammalian system alongside positive and negative expression controls, and after 4 days antibody titers were quantified using bio-layer interferometry with Protein A biosensors. 88.1% of transfectants had detectable antibody titers, ranging from less than 1 µg/ml up to 345 µg/ml (Supplemental Figure 1).

### AlphaLISA specificity binding screen

For accelerated discovery timelines, we developed a highly sensitive bead-based luminescent amplification binding assay using AlphaLISA technology that allows high throughput screening of large antibody panels without need for purification. Unpurified mammalian transfection IgG supernatants were assessed for binding to biotinylated SARS-CoV-2 spike protein, and in parallel for binding to biotinylated irrelevant antigen. Positive and negative control antibodies, (S309 antibody (11) and isotype control antibody, respectively) expressed or spiked in transient transfection supernatants at concentrations matching the range of titers, were included to monitor assay performance. This allowed for fast screening and differentiation between non-binders, non-specific binders and specific binders to the spike protein (Figure 3 and Supplemental Figure 2). Since at this point the supernatants are not concentration normalized, the AlphaLISA signal is a function of both affinity and titer, and thus binders cannot be ranked. Rather, this serves as an efficient process for down-selecting promising candidates that can then be further inquired.

**Figure 3.**
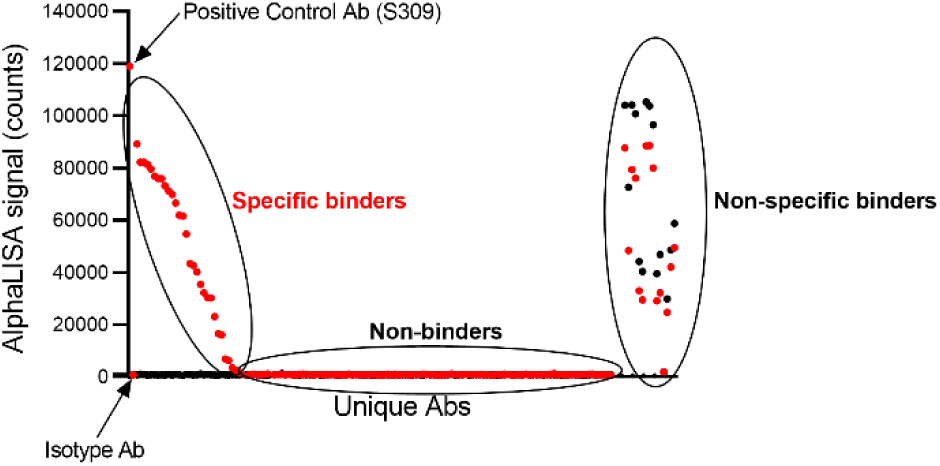
High throughput binding screen allows identification of unique J.HAL^®^ IgG antibody candidates that bind SARS-CoV-2 spike protein. Unpurified transfection supernatants were tested by AlphaLISA for binding to biotinylated SARS-CoV-2 spike protein (red symbols) and in parallel to biotinylated irrelevant target protein (black symbols). Positive control antibody M-3418 (S309) demonstrates binding specificity, whereas isotype control antibody demonstrates lack of binding. Replicates were included when possible. Data was graphed using GraphPad Prism software. Representative binding profile shown.

A total of 73 unique antibody sequences specific for SARS-CoV-2 spike protein were identified: 66 unique antibody sequences identified from the biopanning with Wuhan RBD (20 of which were recovered via NGS mining), and 7 additional unique antibody sequences identified from the biopanning with B.1.1.7 S1 protein. Positive supernatants identified in the initial binding screen were concentration normalized and dose-dependent binding to the SARS-CoV-2 spike protein was assessed by AlphaLISA method described above. (Figure 4 and Supplemental Figure 3). Thus, binding profiles of candidates were evaluated, and candidates were ranked relative to each other and to the positive control S309 antibody. S309 antibody, a high affinity anti-SARS-CoV-2 spike monoclonal antibody (11) displayed dose-dependent binding with a EC_50_ value of 4.2 ng/ml and a Emax value of 121545 (counts) (Supplemental Table 1). All J.HAL^®^ antibodies tested displayed dose-dependent binding, with varying binding potencies (EC_50_ values) and maximum binding (Emax values). Several J.HAL^®^ antibodies display strong binding profile, comparable, albeit inferior to S309 control. For example, antibodies, M-3376, M-3406 and M-3382 display potent binding to the cognate antigen, with EC_50_ values of 8.9, 7.2 and 4.4 ng/ml, respectively and Emax values of 75566, 80909 and 80143 (counts), respectively. This demonstrates the potential of the J.HAL^®^ library to produce specific binders with strong binding profile in absence of affinity maturation.

**Figure 4.**
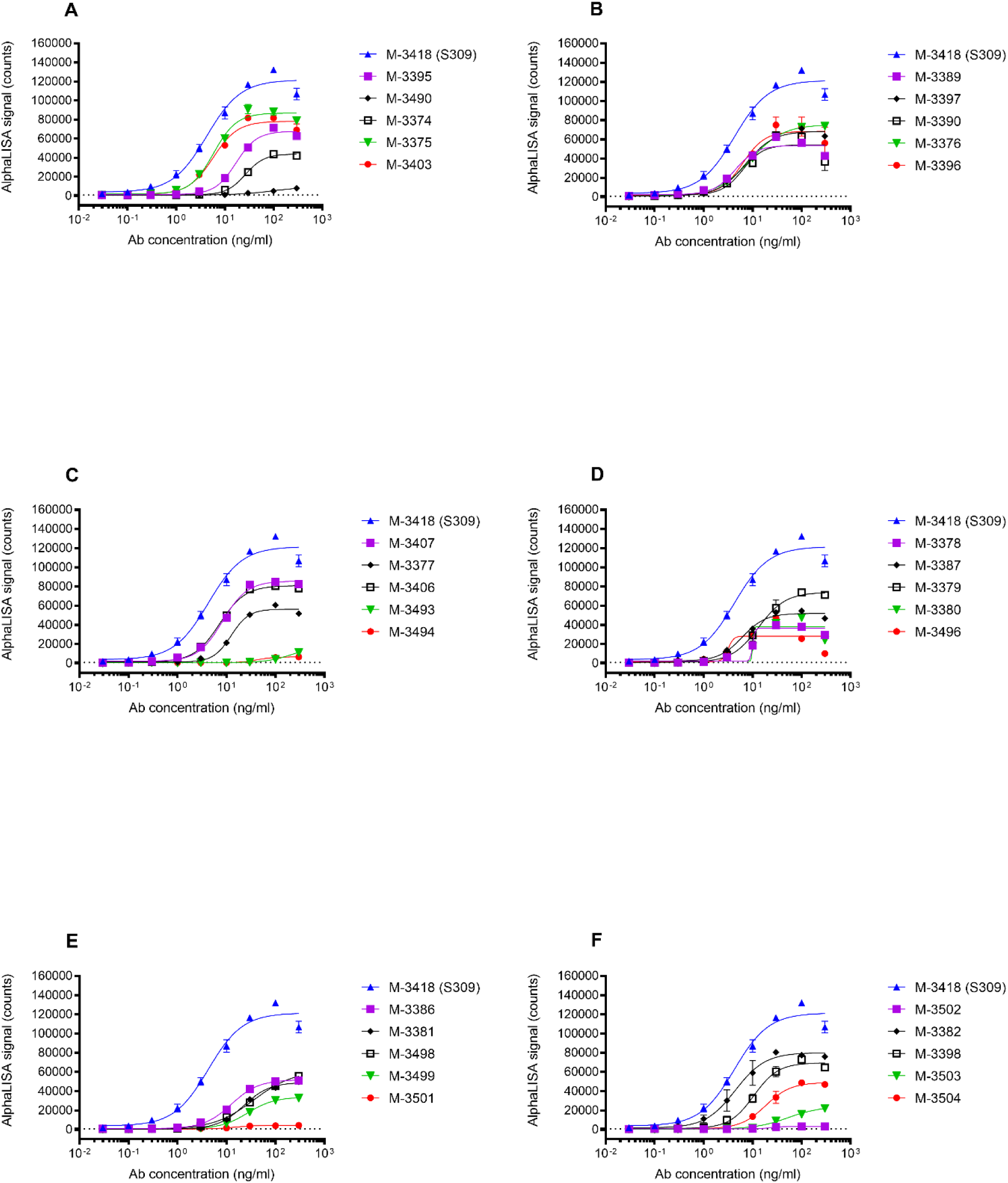
J.HAL^®^ IgG Antibodies exhibit dose-dependent binding to SARS-CoV-2 spike protein. Unpurified transfection supernatants were concentration normalized and tested for binding in a 9-point serial titration series to biotinylated SARS-CoV-2 Spike protein by AlphaLISA. Replicates were included when possible. Data was graphed using GraphPad Prism software. Representative binding profiles for 30 J.HAL^®^ antibodies are shown in the sub-plots A-F above, along with the positive control antibody M-3418 (S309).

### Antibody inhibition screening of spike:human ACE-2 receptor interaction

To quickly identify antibodies that block binding of SARS-CoV-2 spike protein to human ACE-2 receptor, unpurified antibody supernatants that specifically bound SARS-CoV-2 spike protein were tested for their ability to block binding of SARS-CoV-2 spike protein to human ACE-2 by functional ELISA. Antibody blockade, represented as percent inhibition relative to the binding of spike protein alone, was calculated and graphed using GraphPad Prism software. Positive control included 2381 blocking antibody isolated from a convalescent patient (12, 13) spiked at 30 µg/ml in transient transfection supernatant; isotype control antibody, expressed or spiked in transient transfection supernatants at concentrations matching the range of titers, were included to monitor assay performance. Replicates were included when possible and/or data was reproduced in independent experiments. Antibody blockade data is shown in Figure 5 A & B (Mean % Inhibition +/- SD). Since the supernatants are tested neat and are not concentration normalized, the % Inhibition value is a function of both potency and titer, and thus blocking antibodies cannot be ranked, rather selected for further analysis. 36 IgG candidates demonstrated ≥30% blockade and were selected for further analysis. Their respective potencies were further characterized in dose-dependent studies relative to positive benchmark control antibody (Figure 6). A range of potencies were observed, which is as expected. Several IgG candidates demonstrate effective blockade of spike protein binding to human ACE-2 receptor, notably M-3422, M-3388 and M-3406 displaying IC_50_ values of 38 ng/ml, 70 ng/ml and 618 ng/ml, respectively. (Supplemental Table 2).

**Figure 5.**
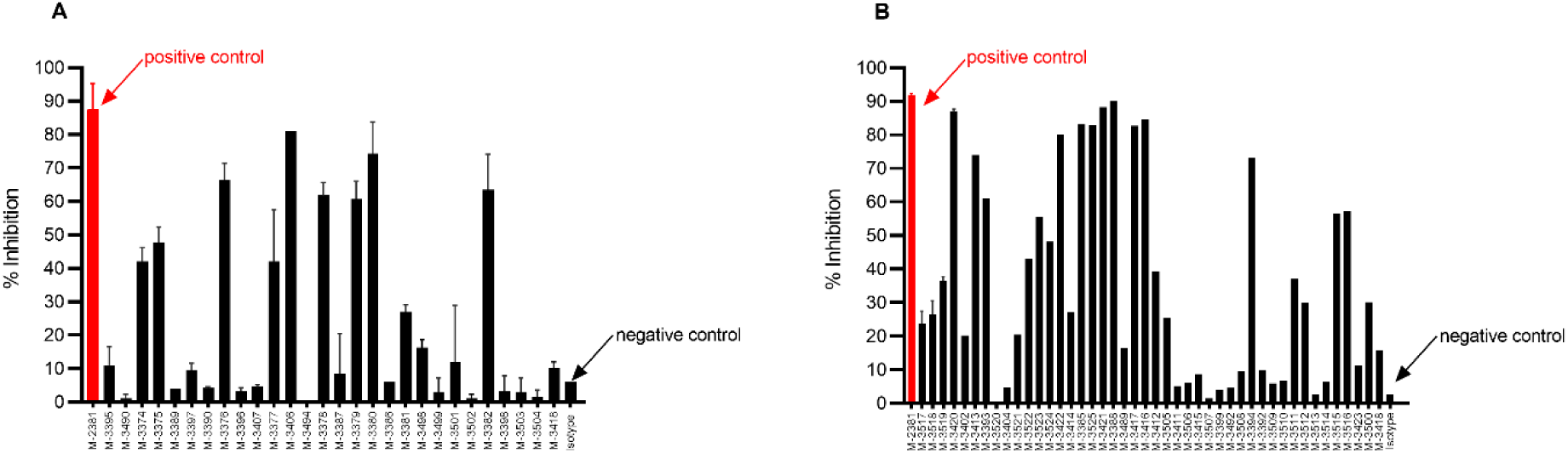
J.HAL^®^ IgG antibodies effectively block Spike:human ACE2 receptor interaction. Functional screen allows identification of IgG antibodies that block SARS-CoV-2 Spike: human ACE-2 receptor interaction. Unpurified transfection supernatants were tested for their ability to block binding of SARS-CoV-2 Spike protein to human ACE-2 using ELISA. Positive control antibody M-2381 (red symbol) demonstrates effective blockade, whereas isotype control antibody demonstrates lack thereof. Replicates were included when possible. Two independent screens were performed (A & B). Data was graphed as % Inhibition +/- SD using GraphPad Prism software.

**Figure 6.**
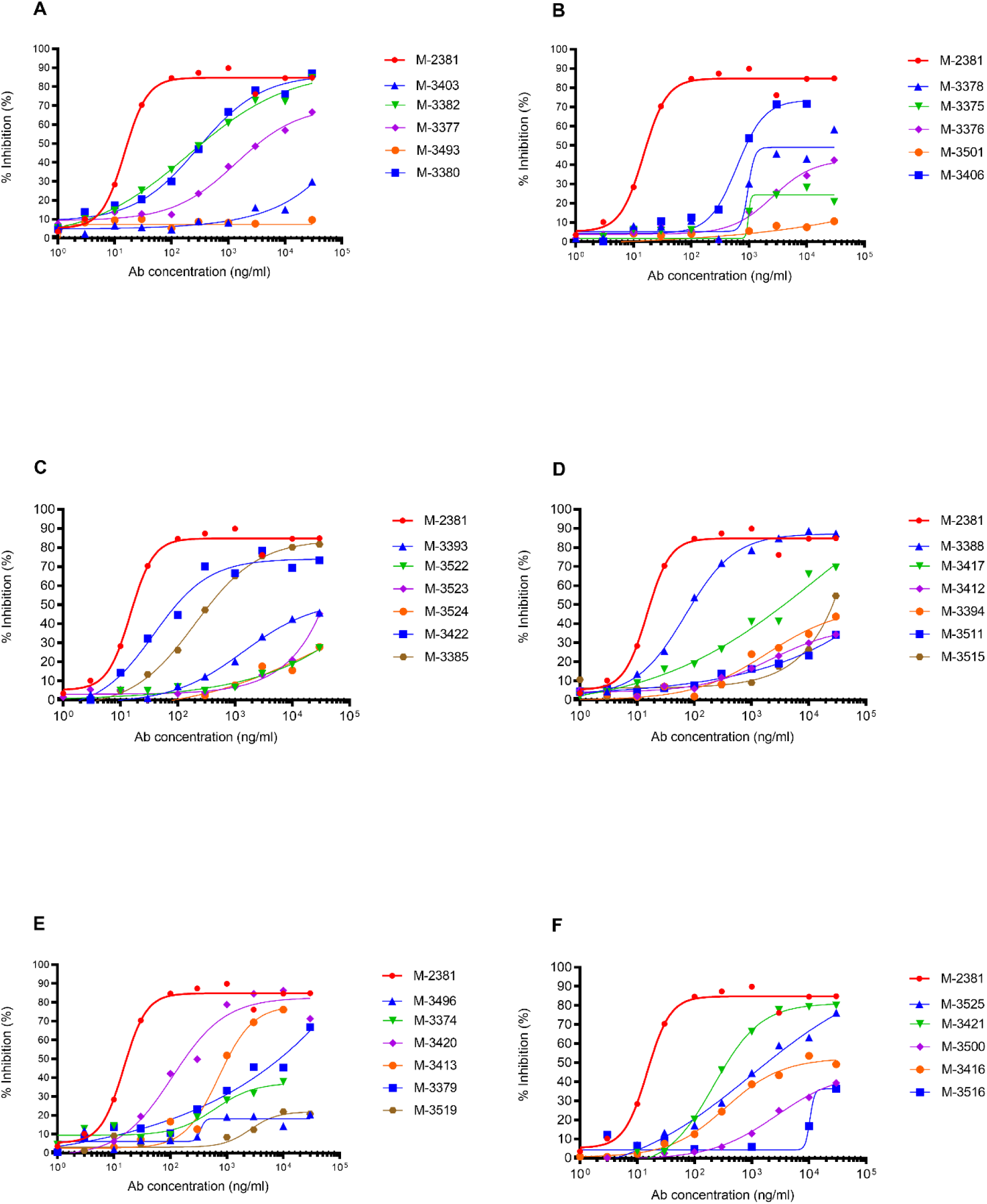
J.HAL^®^ IgG Antibodies exhibit dose-dependent blockade of SARS-CoV-2 Spike: human ACE-2 receptor interaction. Unpurified transfection supernatants were concentration normalized and tested for blocking activity in a 10-point serial titration series. Replicates were included when possible. Data was graphed as % Inhibition +/- SD using GraphPad Prism software. Representative binding profiles for 33 J.HAL^®^ antibodies are shown in the sub-plots A-F above, along with the positive control antibody M-2381 are shown.

### Evaluation of the neutralization activity of IgG candidates using rVSV pseudotyped with the spike of SARS-CoV-2 UK variant (B1.1.7, Alpha)

To evaluate their potency to neutralize SARS-CoV-2 infection, IgG candidates were tested for their ability to neutralize the infection of a rVSV pseudotyped with the spike of the UK variant (B1.1.7, Alpha). For that, two successive screenings were performed in the A549-ACE2/TMPRSS2 cell line. The neutralization activity of IgG candidates was evaluated at 1 µg/ml and 10 µg/ml. Infection data were compared to infected (rVSVΔG-SARS-CoV-2-Spike) and not infected controls. Among all IgG candidates evaluated in the first screen, M-3375, M-3380, M-3403, M-3376, M-3406, M-3412 and M-3420 demonstrated ≥70% of neutralizing activity at 10 µg/ml (Figure 7 A and B). Among these, M-3380 and M-3406 were also capable to inhibit infection by 70% at 1 µg/ml. These 7 IgG candidates were selected for further analysis. No IgG candidates were selected from the second screen (data not shown).

**Figure 7.**
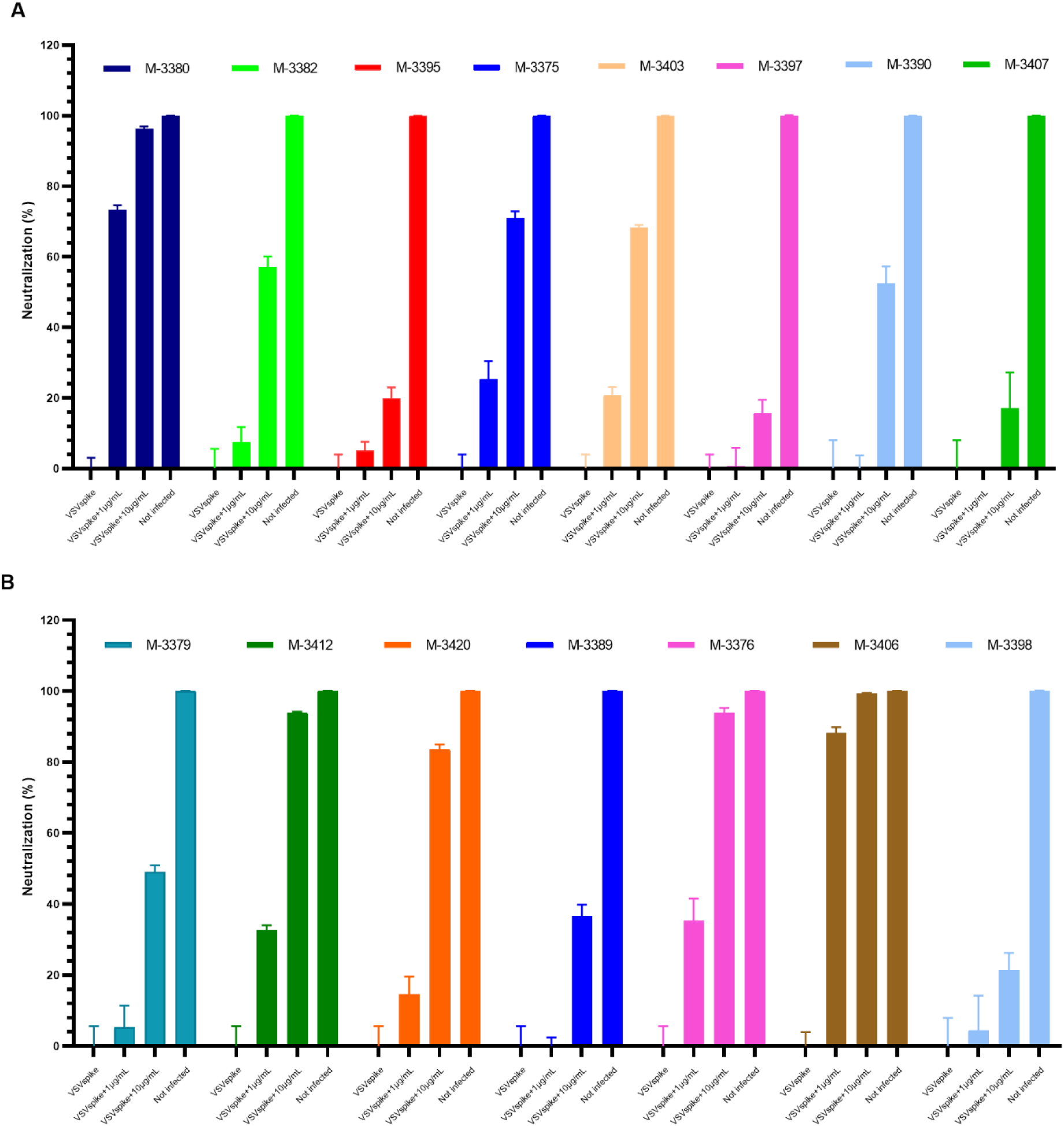
Neutralization activity of J.HAL^®^ IgG antibody candidates (screen 1) on VSV pseudotyped with SARS-CoV-2 UK variant spike (B1.1.7, Alpha) in A549-ACE2/TMPRSS2. IgG candidates were tested for their ability to neutralize the entry of VSV pseudotyped with the spike of SARS-CoV-2 UK variant (B1.1.7, Alpha) in A549-ACE2/TMPRSS2 cells (A: M-3380 to M-3407, B: M-3379 to M-3398). For neutralization experiments, the pseudovirus was pre-incubated with two different concentrations of IgG candidates for 1h at 37°C and the pseudovirus/antibody was subsequently added onto the cells and incubated for 20h at 37°C. Infectivity was assessed at 20 h post-infection by Luciferase assay. The results were normalized to a percentage of neutralization where the cell control (not-infected condition) and the virus control (non-treated, rVSVΔG-SARS-CoV-2-Spike) are used to set 100% neutralization and 0% neutralization, respectively. Data was graphed as of % neutralization + SD using GraphPad Prism software.

### Evaluation of the cross-neutralization activity of selected IgG candidates on a panel of rVSVΔG-SARS-CoV-2-Spike pseudotypes

To determine whether the neutralizing activity of the 8 IgG candidates is conserved across SARS-CoV-2 variants, neutralization assays were performed on a panel of rVSVΔG-SARS-CoV-2-Spike pseudotypes in A549-ACE2/TMPRSS2 cell line. This includes spike proteins from variants of interest and concern: Wuhan (lineage A), D614G mutant (lineage A), Alpha (lineage B.1.1.7), Beta (lineage B.1.351), Gamma (lineage P1), Delta (lineage B.1.617.2), Omicron BA.1 (lineage B.1.1.529) and Omicron BA.2 (lineage B.1.1.529.2). To determine their 50% neutralization dose (ND_50_), defined as the concentration of antibody that reduced the number of infected cells by 50%, increasing concentrations of IgG candidates ranging from 1 ng/ml to 10 µg/ml were used. Table 1 summarizes the ND_50_s calculated for all the IgG candidates. Results indicated that IgG candidates can be classified in 3 groups. In the first group, M-3380 and M-3420 showed a potent neutralization across Wuhan, D614G, Alpha and Delta variants with an ND_50_ between 0.12 and 2.1 µg/ml, but they showed no effect on variants Beta, Gamma and Omicron. In group 2, M-3375, M-3403 and M-3412 showed a decreased neutralization potency against Wuhan and D614G variants compared to Alpha but they were highly potent against Beta and Gamma variants with an ND_50_ between 0.2 and 0.45 µg/ml. As for group 1, they showed no effect on Omicron variant. Finally, the group 3 contains 2 complementary IgG candidates, M-3376 and M-3406. Both showed a potent neutralization across Wuhan, D614G, Alpha and Delta variants but M-3406 was the most potent with an ND_50_ between 0.04 and 0.31 µg/ml. Conversely, only M-3376 was capable to neutralize Beta, Gamma and Omicron variants with an ND_50_ between 0.2 and 0.68 µg/ml. Detailed results obtained for M-3376 and M-3406 are depicted Figure 8. Since M-3376 and M-3406 were highly effective against the most recent variants of concern Delta and Omicron, they were selected for further analysis against SARS-CoV-2 isolates.

**Figure 8.**
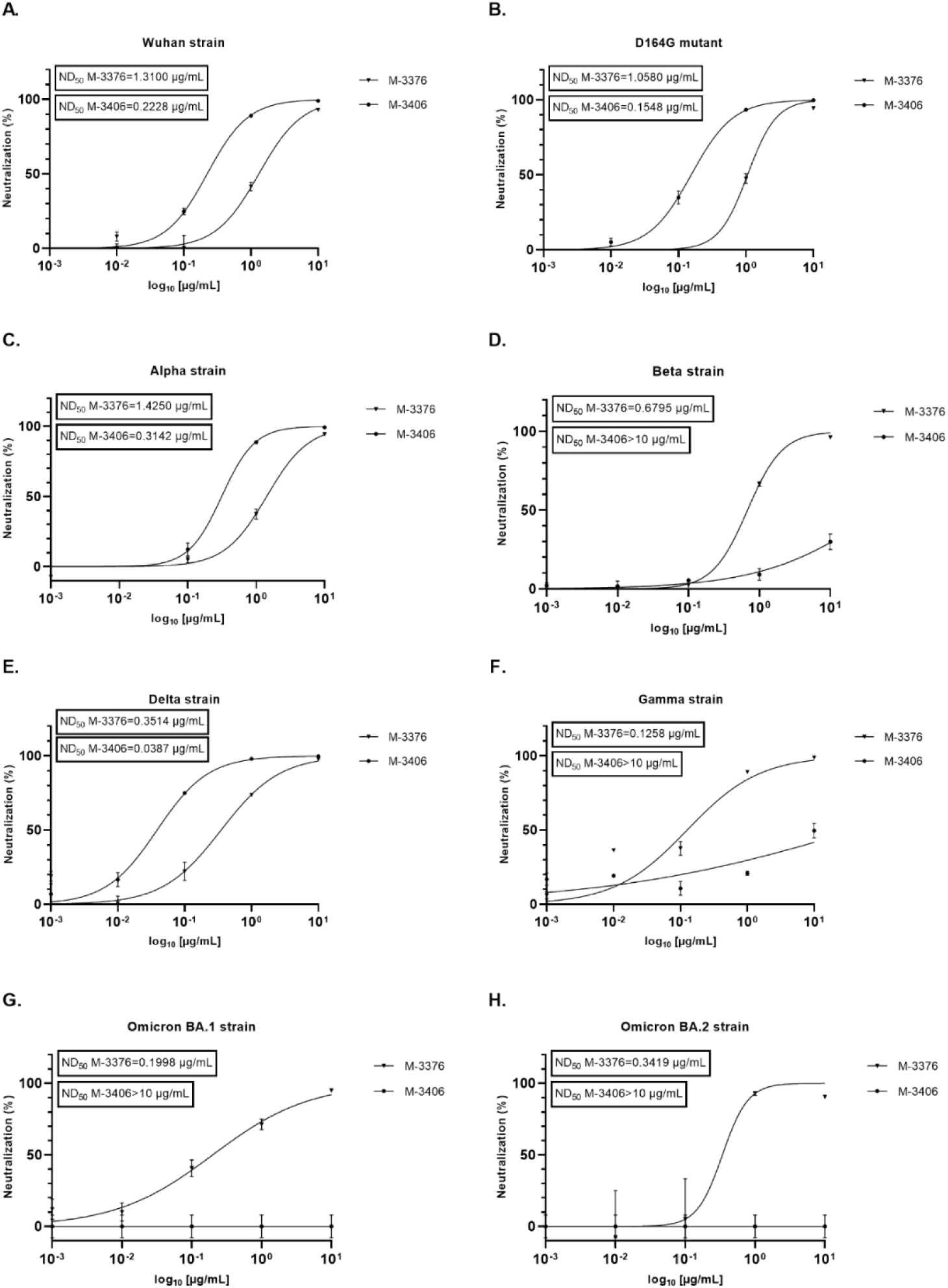
Evaluation of the cross-neutralization activity of J.HAL^®^ IgG candidates M-3376 and M-3406 on a panel of VSV pseudotyped with SARS-CoV-2 spike in A549-ACE2/TMPRSS2 cell line. The neutralizing effects of IgG candidates M-3376 and M-3406 on entry of several rVSVΔG-SARS-CoV-2-Spike pseudotypes was evaluated in A549-ACE2/TMPRSS2 cells. VSV carrying the spike from Wuhan (A), Wuhan D614G mutant (B), Alpha (C), Beta (D), Delta (E), Gamma (F), Omicron BA.1 (G) and Omicron BA.2 (H) were used. For neutralization experiments, the pseudovirus was pre-incubated with a dose range of IgG candidates for 1h at 37°C and the pseudovirus/antibody was subsequently added onto the cells and incubated for 20h at 37°C. Infectivity was assessed at 20 h post-infection by Luciferase assay. The results were normalized to a percentage of neutralization where the cell control (not-infected condition) and the virus control (non-treated, rVSVΔG-SARS-CoV-2-Spike) are used to set 100% neutralization and 0% neutralization, respectively. Data was graphed as of % neutralization + SD and the 50% neutralization dose (ND_50_), defined as the concentration of antibody that reduced the number of infected cells by 50% calculated using GraphPad Prism software.

**Table 1.**
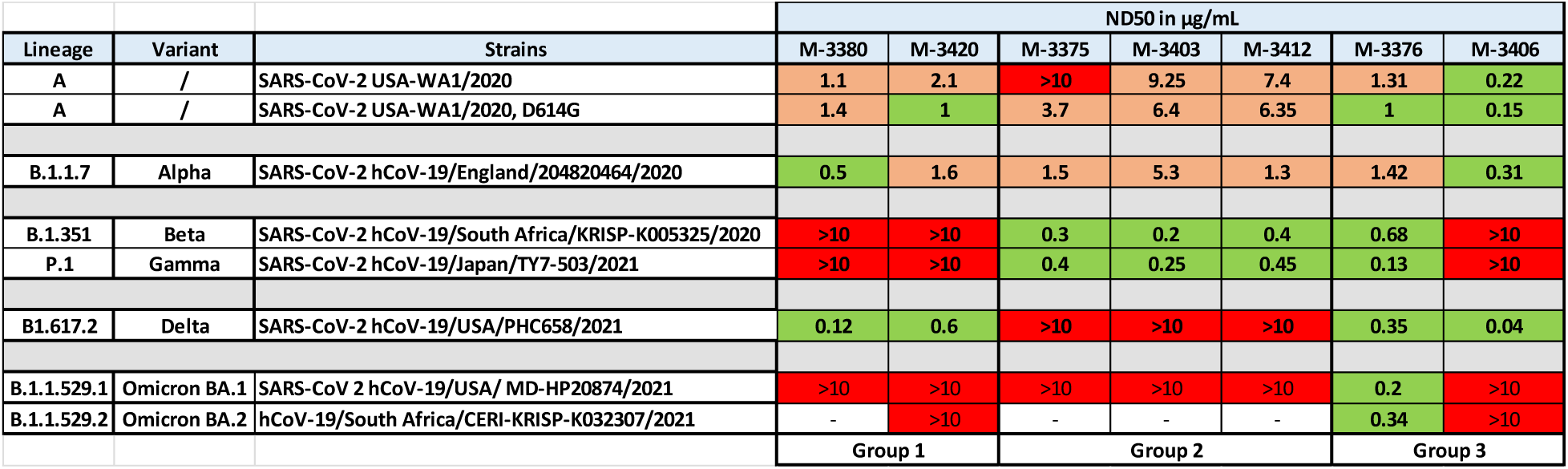
J.HAL^®^ IgG candidates’ neutralization activity on a panel of VSV pseudotyped with SARS-CoV-2 spike in A549-ACE2/TMPRSS2 cell line. The neutralizing effects of IgG candidates on entry of several rVSVΔG-SARS-CoV-2-Spike pseudotypes was evaluated in A549-ACE2/TMPRSS2 cells. VSV carrying the spike from Wuhan, Wuhan D614G mutant, Alpha, Beta, Delta, Gamma, Omicron BA.1 and Omicron BA.2 were used. For neutralization experiments, the pseudovirus was pre-incubated with a dose range of IgG candidates for 1h at 37°C and the pseudovirus/antibody was subsequently added onto the cells and incubated for 20h at 37°C. Infectivity was assessed at 20 h post-infection by Luciferase assay. The 50% neutralization dose (ND_50_), defined as the concentration of antibody that reduced the number of infected cells by 50% calculated using GraphPad Prism software are indicated in the graph. IgG candidates were classified in 3 groups. Green: ND_50_ ≤ 1µg/ml, Orange: 1µg/ml < ND_50_ < 10µg/ml, Red: ND_50_ >10µg/ml.

| Lineage | Variant | Strains | ND50 in µg/mL |  |  |  |  |  |  |
| --- | --- | --- | --- | --- | --- | --- | --- | --- | --- |
|  |  |  | M-3380 | M-3420 | M-3375 | M-3403 | M-3412 | M-3376 | M-3406 |
| A | / | SARS-CoV-2 USA-WA1/2020 | 1.1 | 2.1 | >10 | 9.25 | 7.4 | 1.31 | 0.22 |
| A | / | SARS-CoV-2 USA-WA1/2020, D614G | 1.4 | 1 | 3.7 | 6.4 | 6.35 | 1 | 0.15 |
| B.1.1.7 | Alpha | SARS-CoV-2 hCoV-19/England/204820464/2020 | 0.5 | 1.6 | 1.5 | 5.3 | 1.3 | 1.42 | 0.31 |
| B.1.351 | Beta | SARS-CoV-2 hCoV-19/South Africa/KRISP-K005325/2020 | >10 | >10 | 0.3 | 0.2 | 0.4 | 0.68 | >10 |
| P.1 | Gamma | SARS-CoV-2 hCoV-19/Japan/TY7-503/2021 | >10 | >10 | 0.4 | 0.25 | 0.45 | 0.13 | >10 |
| B1.617.2 | Delta | SARS-CoV-2 hCoV-19/USA/PHC658/2021 | 0.12 | 0.6 | >10 | >10 | >10 | 0.35 | 0.04 |
| B.1.1.529.1 | Omicron BA.1 | SARS-CoV 2 hCoV-19/USA/ MD-HP20874/2021 | >10 | >10 | >10 | >10 | >10 | 0.2 | >10 |
| B.1.1.529.2 | Omicron BA.2 | hCoV-19/South Africa/CERI-KRISP-K032307/2021 | - | >10 | - | - | - | 0.34 | >10 |
|  |  |  | Group 1 |  | Group 2 |  |  | Group 3 |  |

### Confirmation of the cross-neutralization activity of IgG candidates M-3376 and M-3406 on a panel of SARS-CoV-2 isolates

To validate their potency, the capacity of IgG candidates M-3376 and M-3406 to neutralize SARS-CoV-2 isolates, WA.1, Delta and Omicron BA.1 was evaluated in A549-ACE2/TMPRSS2 cell line. A negative control antibody was used in parallel as well as 2 positive controls, the S309 neutralizing antibody (11) and remdesivir, a nucleoside analogue inhibiting SARS-CoV-2 replication (14). Neutralization was performed by incubating increasing concentrations of IgG candidates ranging from 0.01 µg/ml to 100 µg/ml with the virus at MOI 0.1 for 1 hour prior to infection of cells. Remdesivir was added 1hour post-infection. Viral titers (expressed as genome equivalent per µl of RNA) were quantified in cell lysates by RT-qPCR 40 hours post-treatment and results of treated conditions were compared to the infected control. Results indicated that contrary to the negative control antibody, S309 was effective against the 3 isolates (Figure 9). Indeed, the viral titers decreased in a dose dependent-manner to reach 1.5-log reduction at the highest concentration tested (100 µg/ml). Remdesivir also reduced viral replication of WA.1, Delta and Omicron BA.1 isolates by 2-log, 2.5-log and 3-log respectively. A slightly higher reduction level (2.5-log) was observed against WA.1 isolate using M-3376 and M-3406 at the highest concentration (100 µg/ml) (Figure 9A). However, no antiviral effect was observed at the other concentrations tested. These two IgG candidates were more potent than remdesivir and S309 against the Delta isolate with a 2.5-3-log reduction observed for M-3376 at 10 µg/ml and for M-3406 at 1 µg/ml (Figure 9B). However, only M-3376 conserved an antiviral activity comparable to remdesivir and superior to S309 against Omicron BA.1 with a 1.5-log and 3-log reduction using 10 µg/ml and 100 µg/ml respectively (Figure 9C). Altogether, these results indicate that M-3376 and M-3406 were able to neutralize SARS-CoV-2 isolates. They also confirm their complementarity since M-3376 neutralized Omicron BA.1 isolate and M-3406 was superior to neutralize Delta isolate.

**Figure 9.**
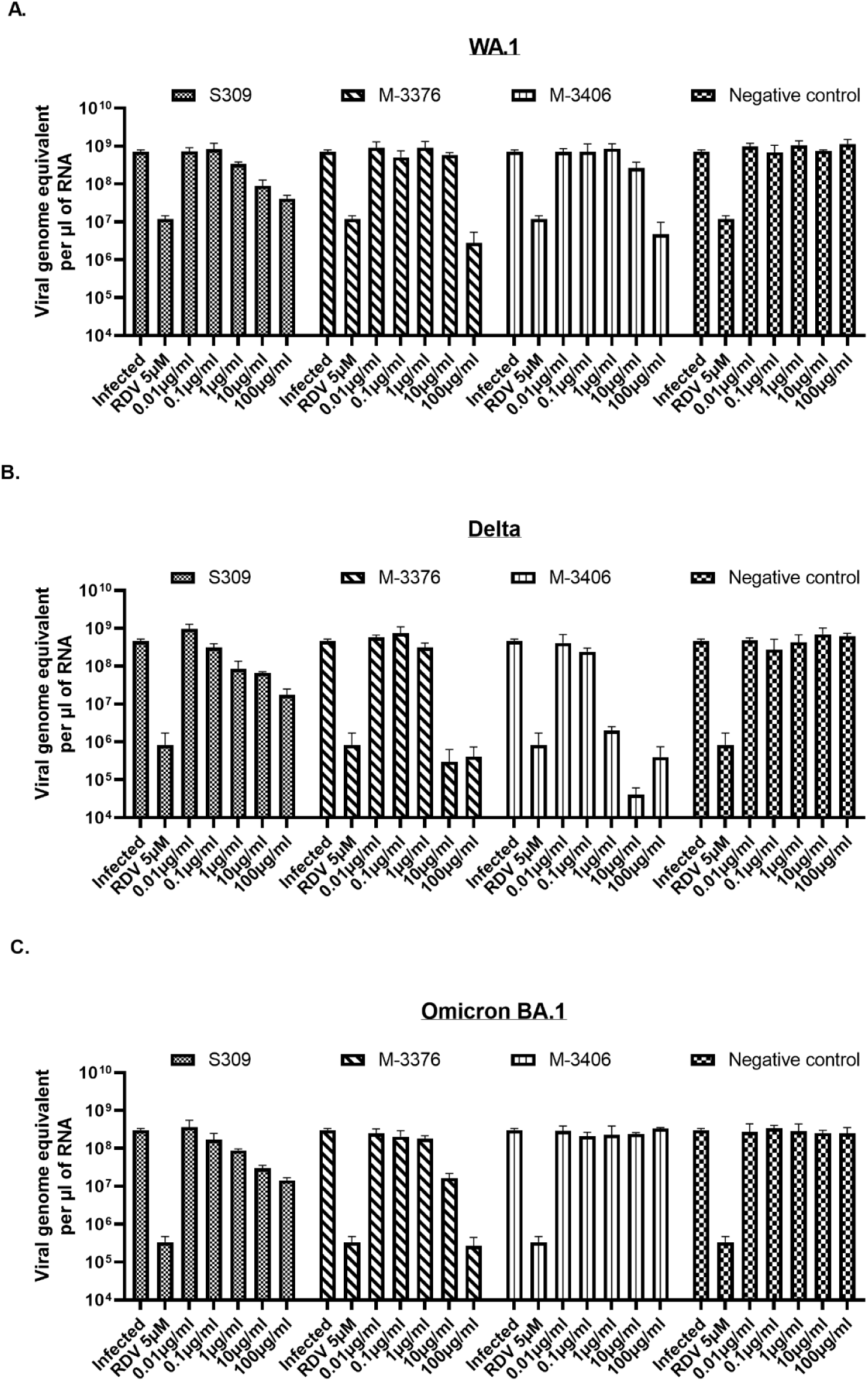
Confirmation of the cross-neutralization activity of IgG candidates M-3376 and M-3406 on a panel of SARS-CoV-2 isolates in A549-ACE2/TMPRSS2 cell line. The neutralizing effects of IgG candidates M-3376 and M-3406 on entry of SARS-CoV-2 isolates WA.1 (A), Delta (B) and Omicron BA.1 (C) was evaluated in A549-ACE2/TMPRSS2 cells. For neutralization experiments, the virus at MOI 0.1 was pre-incubated with a dose range of IgG candidates for 1h at 37°C and the virus/antibody was subsequently added onto the cells. Positive (S309) and negative control antibodies were also subjected to the same protocol. Remdesivir, a nucleoside analogue, was added 1h post-infection and used as a second positive control. Viral titers were quantified in cell lysates by RT-qPCR 40hrs post-treatment using the standard curve method. Data was graphed as viral titers (expressed in genome equivalent per µl of RNA) + SD using GraphPad Prism software.

### Kinetic binding affinity assessment

To further assess the target specific binding profile of the lead candidates, kinetic binding profiles of the lead candidates against 8 clinically relevant strains were assessed by SPR using the Carterra LSA instrument (carterra-bio.com). All kinetics measurements were performed at 25°C using both purified and unpurified IgG candidates. Many of the J.HAL^®^ lead candidates displayed respectable target binding affinity (k_D_ 1-10 nM range) and cross-reactivity against spike target antigens from several viral strains of interest (Figure 10). Additionally, we observed good correlation between the SPR binding profiles of purified candidates and unpurified supernatants (data not shown), validating the feasibility to use SPR technology to assess pan cross-reactivity of unpurified transfection supernatants to accelerate discovery timelines.

**Figure 10.**
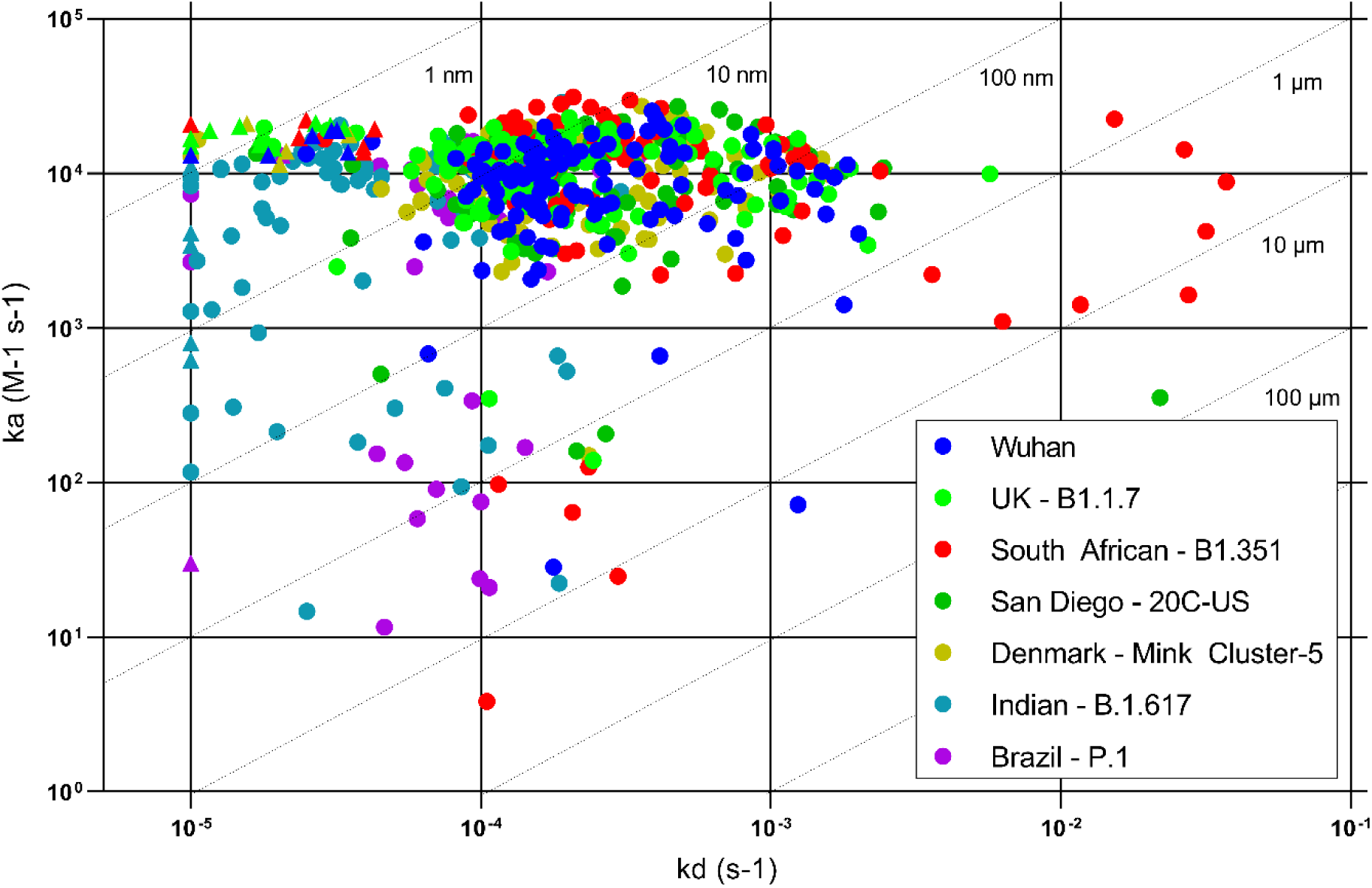
SPR demonstrating cross-reactivity of top J.HAL^®^ antibody candidates. Triangle markers are the S309 control antibody. All other markers on the iso-affinity plot represent one J.HAL^®^ IgG binding interaction with one of the seven SARS-CoV-2 strains as indicated in the legend.

### Biophysical characterization

Figure 11 shows the behavior of the selected J.HAL^®^ antibodies across 6 developability assays in our platform. Conformational stability was assessed by differential scanning fluorimetry (DSF), low pH hold, and chemical unfolding. Colloidal stability was assessed by polyethylene glycol (PEG) solubility, self-interaction nanoparticle spectroscopy (SINS), and standup monolayer affinity chromatography (SMAC). DSF assesses the temperature at which certain regions of the antibody begin to unfold. More stable, and thus more developable antibodies tend to have one or more regions unfold at higher temperatures and have a higher first-unfolding transition temperature. The first transition (T1) correlates with the CH2 domain, and the second transition (T2) correlates with the unfolding of the Fab and CH3 domain regions. Lack of a second transition temperature is indicative of the Fab unfolding at the same or similar temperature to the CH2 domain which is suggestive of lower stability. The DSF data in Figure 11A is reported as weighted shoulder score (WSS) which is a metric for thermal stability that can be derived from a DSF spectrum. The WSS algorithm consists of identifying the T1 peak, normalizing the spectrum to the intensity of the T1 peak, and integrating the area under the curve between the T1 and the temperature where the intensity drops below the baseline. The integration is weighted by the squared temperature difference from T1. A higher WSS value indicates multiple transition temperatures and therefore improved stability, with WSS values increasing as the second transition temperature increases. Many of the J.HAL^®^ lead candidates showed high WSS indicating suitable thermal stability.

**Figure 11.**
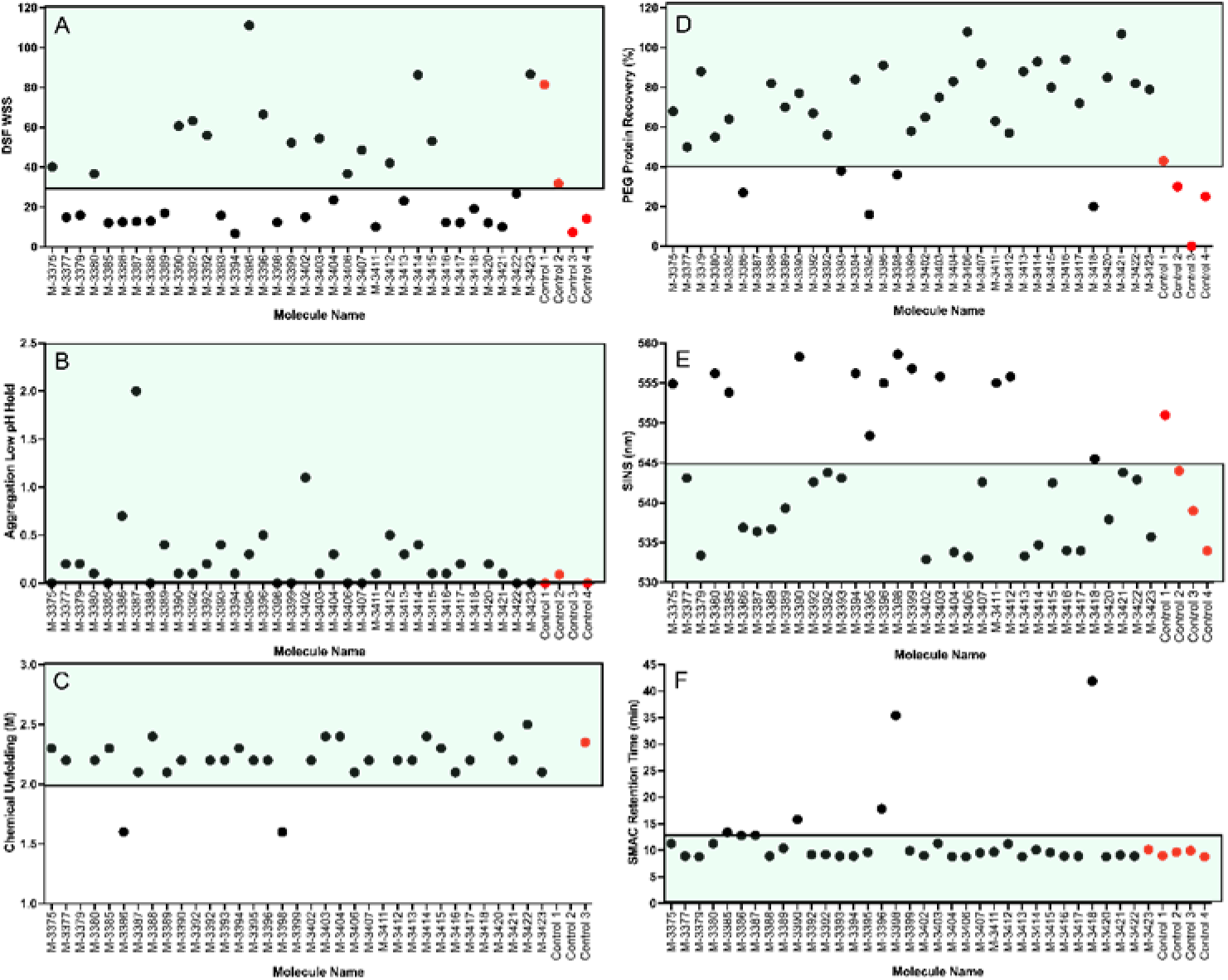
Antibody biophysical characterization indicating potential for manufacturability. For respective LC and HC amino acid sequences of these homodimeric antibodies, listed here by name and Molecule ID number. Conformational stability results are shown in panels **A.** DSF weighted shoulder score (WSS), **B.** aggregation after low pH hold and **C.** chemical unfolding. Colloidal stability results are shown in panels **D.** relative PEG solubility, **E.** SINS wavelength maximum and **F.** SMAC retention time. Acceptable value range for favorable developability characteristics is shown in green and system suitability standards are indicated in red symbols

Protein-protein self-interaction has been correlated to low solubility, high viscosity and increased manufacturability difficulties. Therefore, it is helpful to understand and screen out molecules early in development that self-interact. The SINS assay utilizes gold colloid surfaces to measure protein-protein interaction. Gold colloids have unique optical absorption properties dependent on their aggregation state, absorbing light at longer wavelengths as nanoparticle aggregation occurs. Protein self-interaction is assessed by capturing test antibodies on the surface of a gold colloid and measuring the shift of the absorbance maxima. If the immobilized antibodies self-interact, the absorbance maximum of the spectrum red shifts to longer wavelengths (545-565 nm), or blue shifts to shorter wavelengths (530-545 nm), if less interaction occurs. The J.HAL^®^ candidates displayed a wide variety of SINS results, with many of the candidates showing short wavelengths indicating no protein-protein self-interaction.

The inherent solubility of a molecule is thought to be related to aggregation levels throughout manufacturing process development and during long-term stability. Solubility was assessed by exposing the protein to PEG which precipitates the protein out of solution. After incubation and filtration of insoluble precipitates, protein concentration is measured, and recovery calculated. Molecules with higher solubility will show more soluble protein recovered after exposure to PEG. Most of the J.HAL^®^ candidates showed well over 50% protein recovery after exposure to PEG solutions, suggesting inherently high solubility.

The SMAC method is used to assess colloidal stability of a molecule by monitoring the retention of a molecule on a Zenix HPLC column. A commercially available Zenix^®^ column (Millipore Sigma) with a hydrophobic standup monolayer can be used; it is hypothesized that molecules with colloidal instability interact with the column resulting in a delayed retention time. Undiluted samples are loaded onto the Zenix^®^ column and are eluted isocratically with a 100 mM sodium phosphate, pH 7.0 running buffer, and absorbance at 220 nm wavelength is monitored. Very few of the J.HAL^®^ candidates interacted with the Zenix^®^ column indicating that the lead candidates have favorable colloidal stability as measured by this method.

The inflection point of a chemical unfolding curve is thought to be related to conformational stability and to stability during long-term storage, with a greater inflection point relating to a more structurally or conformationally stable molecule. The chemical unfolding curve is produced by exposing the molecule to increasing concentrations of the denaturant, guanidine hydrochloride. After 24 hours, the intrinsic fluorescence of the samples was measured using a SUPR-UV plate reader (Protein Stable). The collected raw data were then processed, and the chemical unfolding curve and its inflection point was calculated from the processed data as a function of denaturant condition.

Exposure to low pH during viral inactivation could result in significant unfolding and aggregation in a conformationally unstable molecule. Aggregation that results from low pH instability is cleared by subsequent purification steps but can negatively impact final yield. Stability during low pH viral inactivation was assessed by titration with acetic acid to pH 3.3, held for 30 minutes, and then titrated to a final pH of 5.0. Aggregation was measured by SEC analysis and compared with a control sample that was not exposed to low pH stress. An increase in aggregation of more than 10% indicates the molecule is low pH unstable. None of the J.HAL^®^ candidates showed an increase in aggregate of more than 2 % and are therefore all stable when exposed to pH 3.3.

Altogether, these results demonstrate the potential of our AI-derived library and our high throughput screening approach to identify panels of antibodies with good binding profile and activity without need of affinity maturation. Furthermore, these humanoid antibodies exhibit desirable developability features.

## DISCUSSION

We describe here a novel, cost-effective and accelerated approach to therapeutic antibody discovery, that couples *de novo* human antibodies derived *in silico*, which we refer to as “humanoid antibodies”, with high throughput screening technologies. We have developed the Antibody-GAN, a new in silico engineering approach to designing a novel class of diverse antibody therapeutics that mimic somatically hypermutated human repertoire response. The platform allows antibody biasing towards desirable developability attributes, such as suitability for common protein manufacturing processes, high stability during long-term storage conditions, low viscosity for injectability at high concentration and long elimination half-lives to reduce dosing frequency (3). Here, we employed the Antibody-GAN to design and construct a 1 billion theoretical diversity phage Fab display library. This library is in process of expanding to over 50 billion theoretical diversity by the end of 2023. The first validation campaign described herein was conducted with the intent to discover antibodies to the SARS-CoV-2 spike protein and demonstrate library utility.

Phage display methodology has proven to be a robust, and efficient platform to discover and develop human therapeutic antibodies (15). Our Antibody-GAN generated phage library is comprised of 25 sub-libraries, 5 heavy chain germlines paired combinatorially with 5 variable chain (4 kappa and 1 lambda) germlines. Fv sequence diversity encompasses both framework and CDR regions which drives affinity, efficacy and developability (3). Most synthetic libraries focus only on CDR diversity. Due to the sensitive balance between affinity, efficacy and developability, it is likely advantageous to consider the entire Fv sequence space when engineering the optimal therapeutic candidate. As data is collected on the sequence-function/stability relationship, further library refinement will be conducted, improving discovery and developability success.

The applicability of the humanoid library for effective therapeutic antibody discovery is demonstrated here with the identification of a panel of human monoclonal antibodies that are novel, diverse, and pharmacologically active. These first-generation antibodies, without the need for affinity maturation, bind to the SARS-CoV-2 spike protein with therapeutically relevant specificity and affinity and display efficient inhibition of spike:human ACE2 receptor binding. The binding affinity exhibited has proven sufficient for effective functional activity, first demonstrated through the blockade of spike:ACE2 interaction, then confirmed in neutralization of SARS-CoV-2 viral infectivity across several strains.

Noteworthy, the antibody panel identified encompasses diverse antibodies with different selectivity and potency. Indeed, a group of antibodies, such as M-3380 and M-3420 show potent neutralization across Wuhan, D614G, Alpha and Delta variants, but no effect on variants Beta, Gamma and Omicron. In contrast, a second group of antibodies, such as M-3375, M-3403 and M-3412 show a decreased neutralization potency against Wuhan and D614G variants compared to Alpha but highly potent neutralization against Beta and Gamma variants. Of particular interest are two antibody candidates, M-3376 and M-3406, which display potent and complementary neutralization activity across strains. Both show potent neutralization across Wuhan, D614G, Alpha and Delta variants, although M-3406 is the most potent. Conversely, only M-3376 is able to neutralize Beta, Gamma and Omicron variants. Combined, these two antibody candidates represent an attractive therapeutic option, either co-administered in cocktail form or engineered as a bi-specific therapeutic modality, effective across all SARS-CoV-2 strains. Altogether, this demonstrates the diversity of our humanoid antibody library J.HAL^®^ and the potential of identifying broadly neutralizing antibodies with desirable developability attributes.

Expedient antibody discovery is critical for pandemic response. This platform explicated the isolation, characterization and identification of functional IgG candidates in less than three months. The viral neutralization assays exhibited good correlation with the rVSVΔG-SARS-CoV-2-Spike pseudotypes and SARS-CoV-2 isolates. These observations highlight the robustness of the VSV pseudotyped model for the rapid identification of effective neutralizing antibody candidates.

Furthermore, the SARS-CoV-2 J.HAL^®^ antibodies are anticipated to fit our current process development and manufacturing platform. As such, this would benefit cost-of-goods and improve therapeutic access to patients. Together with the bias towards improved developability attributes and improved in-use characteristics such as low viscosity for injectability at high concentration, the GAN-derived antibodies represent an advantageous therapeutic modality. A modality that can be developed and deployed faster than antimicrobials and vaccines, thus well suited for rapid response to infectious threats, such as pandemic response.

## ACKNOWLEDGEMENTS

We gratefully thank Alex Taylor for bioinformatical support, Kathryn McLean and Lindsay Pautsch for generation and purification of material.

## SUPPLEMENTAL

**Figure S1.**
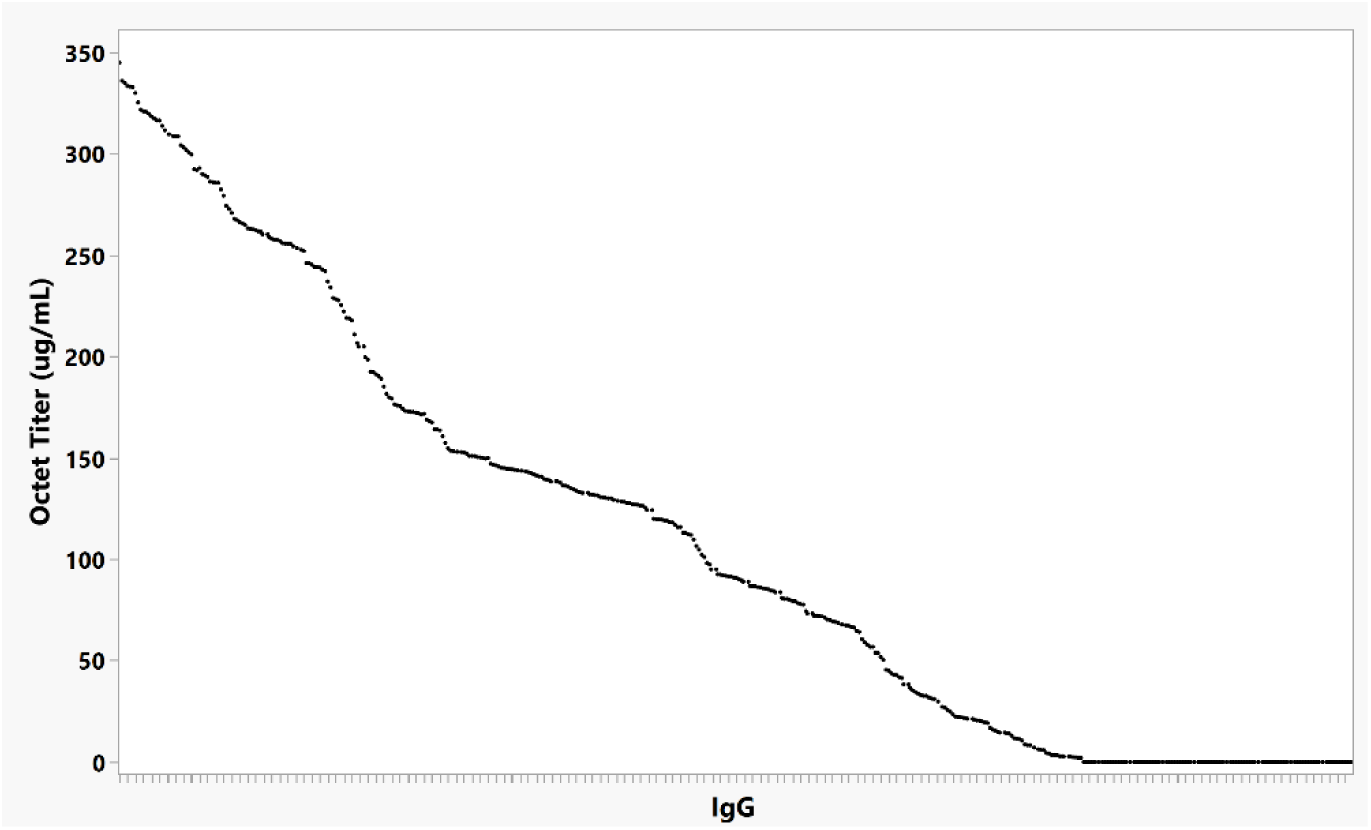
Expi293F titers of 463 IgG sequences after 4 days of expression, as measured by bio-layer interferometry using the Octet® RED96.

**Figure S2.**
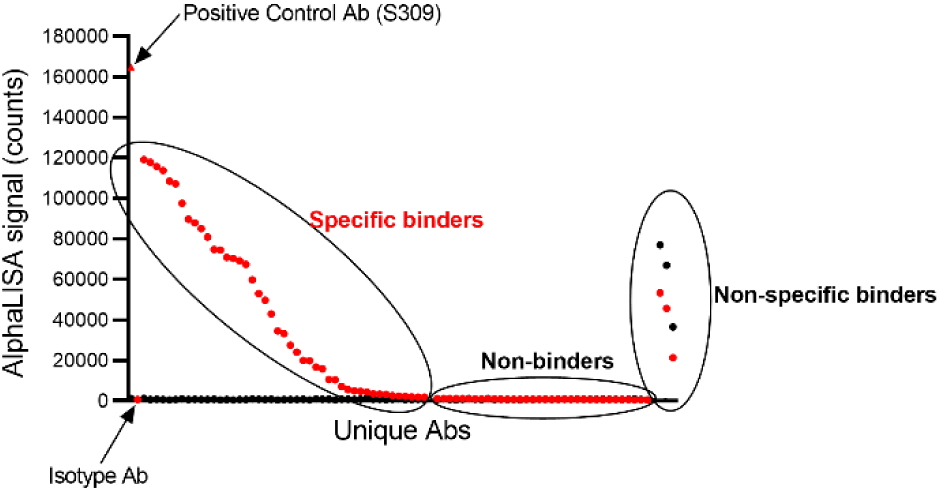
Identification and characterization of unique IgG antibody candidates that bind SARS-CoV-2 spike protein. Binding screen allows identification of target-specific binders. Unpurified transfection supernatants were tested by AlphaLISA for binding to biotinylated SARS-CoV-2 Spike protein (red symbols) and in parallel to biotinylated irrelevant target protein (black symbols). Positive control antibody S309 demonstrates binding specificity, whereas isotype control antibody demonstrates lack of binding. to Replicates were included when possible. Data was graphed using GraphPad Prism software. Representative binding profiles are shown.

**Figure S3.**
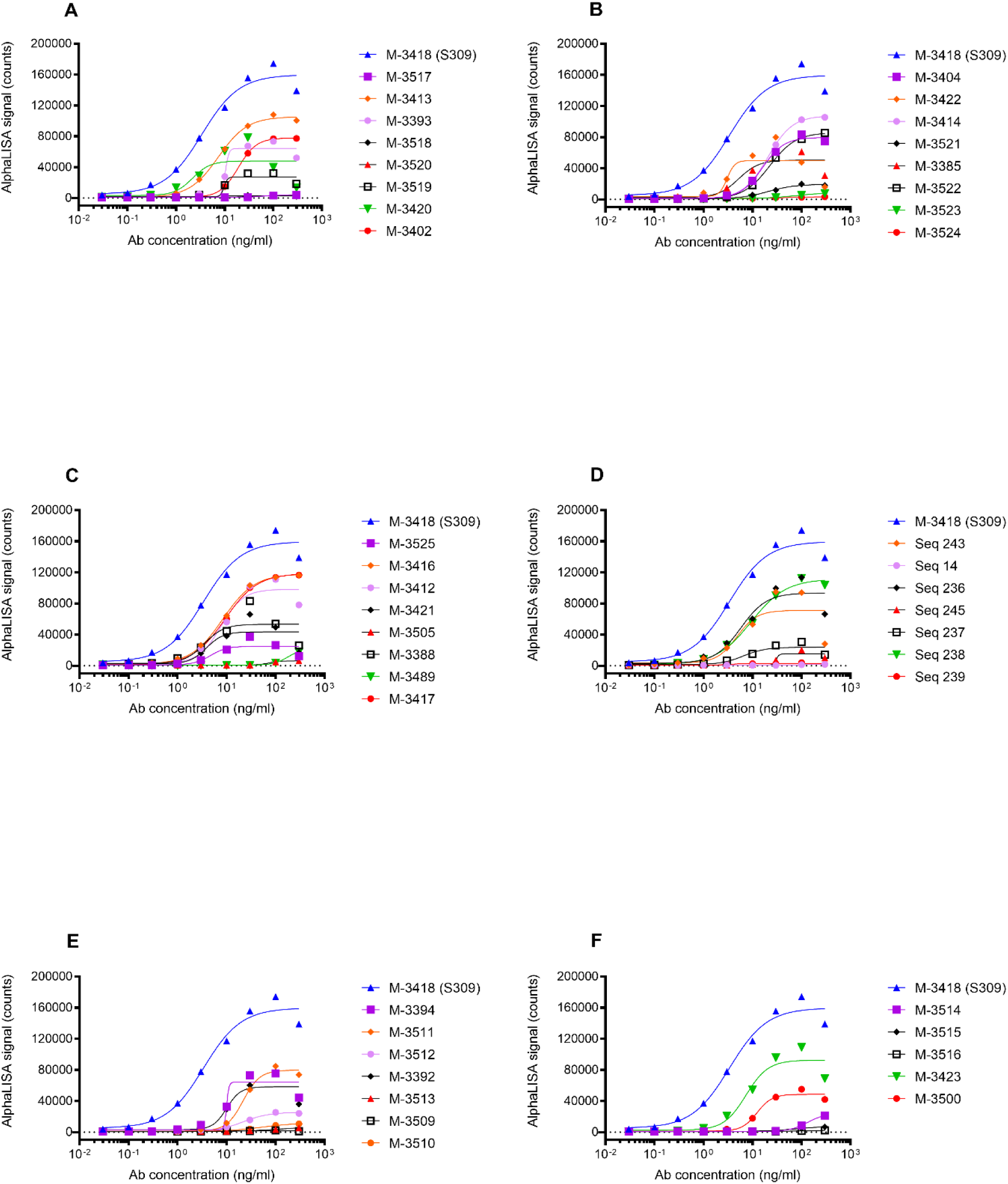
J.HAL^®^ IgG Antibodies exhibit dose-dependent binding to SARS-CoV-2 spike protein. Unpurified transfection supernatants were concentration normalized and tested for binding in a 9-point serial titration series to biotinylated SARS-CoV-2 Spike protein by AlphaLISA. Replicates were included when possible. Data was graphed using GraphPad Prism software. Representative binding profiles for 43 J.HAL^®^ antibodies are shown in the sub-plots A-F above, along with the positive control antibody M-3418 (S309).

**Table S1.** EC_50_ values and Emax values of J.HAL^®^ IgG Antibodies dose-dependent binding to SARS-CoV-2 spike protein.

| **Ab** | **EC<sub>50</sub>**<br>**(ng/ml)** | **E<sub>max</sub>**<br>**value** |
| --- | --- | --- |
| M-3418 | 4.2 | 121545 |
| M-3395 | 17.0 | 67708 |
| M-3490 | 111.7 | 10407 |
| M-3374 | 26.5 | 43875 |
| M-3375 | 5.8 | 86820 |
| M-3403 | 5.7 | 78177 |
| M-3389 | 4.4 | 53712 |
| M-3397 | 7.3 | 68554 |
| M-3390 | 6.3 | 54307 |
| M-3376 | 8.9 | 75566 |
| M-3396 | 6.3 | 68423 |
| M-3407 | 8.3 | 85746 |
| M-3377 | 12.5 | 56439 |
| M-3406 | 7.2 | 80909 |
| M-3493 | 208.3 | 17156 |
| M-3494 | 42.2 | 6682 |
| M-3378 | n/a | 36011 |
| M-3387 | 6.1 | 52083 |
| M-3379 | 13.3 | 73976 |
| M-3380 | n/a | 38133 |
| M-3496 | 3.2 | 28382 |
| M-3386 | 12.6 | 51958 |
| M-3381 | 22.4 | 49261 |
| M-3498 | 36.0 | 61444 |
| M-3499 | 27.2 | 33902 |
| M-3501 | 14.4 | 4319 |
| M-3502 | 23.1 | 3292 |
| M-3382 | 4.4 | 80143 |
| M-3398 | 10.6 | 69524 |
| M-3503 | 52.6 | 24187 |
| M-3504 | 18.5 | 49166 |

**Table S2.** IC_50_ values of J.HAL^®^ IgG Antibodies dose-dependent blocking to SARS-CoV-2 spike protein binding to human ACE-2 receptor protein.

| Ab | IC<sub>50</sub><br>(ng/ml) |
| --- | --- |
| M-2562 | 15 |
| M-3374 | 613 |
| M-3375 | ~ 978.8 |
| M-3403 | N.C. |
| M-3376 | 2574 |
| M-3377 | 1364 |
| M-3406 | 618 |
| M-3493 | N.C. |
| M-3378 | 964 |
| M-3379 | N.C. |
| M-3380 | 293 |
| M-3496 | ~ 342.1 |
| M-3500 | 2732 |
| M-3501 | N.C. |
| M-3382 | 200 |
| M-3412 | 2215 |
| M-3394 | 1892 |
| M-3511 | N.C. |
| M-3515 | N.C. |
| M-3516 | N.C. |
| M-3519 | 2608 |
| M-3420 | 109 |
| M-3413 | 721 |
| M-3393 | 1408 |
| M-3522 | N.C. |
| M-3523 | N.C. |
| M-3524 | N.C. |
| M-3422 | 38 |
| M-3385 | 203 |
| M-3525 | 997 |
| M-3421 | 217 |
| M-3388 | 70 |
| M-3417 | N.C. |
| M-3416 | 353 |

N.C. = not calculated, due to poor sigmoidal curve fit.

